# Wireless Programmable Recording and Stimulation of Deep Brain Activity in Freely Moving Humans

**DOI:** 10.1101/2020.02.12.946434

**Authors:** Uros Topalovic, Zahra M. Aghajan, Diane Villaroman, Sonja Hiller, Leonardo Christov-Moore, Tyler J. Wishard, Matthias Stangl, Nicholas R. Hasulak, Cory Inman, Tony A. Fields, Dawn Eliashiv, Itzhak Fried, Nanthia Suthana

**Affiliations:** Department of Electrical and Computer Engineering, UCLA, Los Angeles, CA 90095, United States; Department of Psychiatry and Biobehavioral Sciences, Jane and Terry Semel Institute for Neuroscience and Human Behavior, UCLA, Los Angeles, CA 90024, United States; NeuroPace Inc., Mountain View, CA 94043, United States; Department of Neurology, UCLA, Los Angeles, CA 90095, United States; Department of Bioengineering, UCLA, Los Angeles, CA 90095, United States; Department of Neurosurgery, David Geffen School of Medicine, UCLA, Los Angeles, CA 90095, United States; Functional Neurosurgery Unit, Tel Aviv Medical Center and Sackler School of Medicine, Tel Aviv University, Tel Aviv, Israel

**Keywords:** Neuroimaging Methods, Human, Intracranial EEG, Deep Brain Stimulation, Wearables, Virtual Reality, Augmented Reality

## Abstract

Current implantable devices that allow for recording and stimulation of brain activity in humans are not inherently designed for research and thus lack programmable control and integration with wearable sensors. We developed a platform that enables wireless and programmable intracranial electroencephalographic recording and deep brain stimulation integrated with wearable technologies. This methodology, when used in freely moving humans with implanted neural devices, can provide an ecologically valid environment conducive to elucidating the neural mechanisms underlying naturalistic behaviors and developing viable therapies for neurologic and psychiatric disorders.

## Main

Traditional methods for recording and modulating brain activity in humans (e.g., fMRI, MEG, TMS) require immobility and are thus limited in their application during laboratory-based tasks low in ecological validity. Given the recent increase in medical therapies using implanted neural devices to treat and evaluate abnormal brain activity in patients with epilepsy [1] and other neurologic and psychiatric disorders [2-6], recording and modulating deep brain activity during freely moving behavior is now possible. There are already over two thousand individuals with chronic sensing and stimulation electrodes implanted within the brain, with the number expected to increase as additional invasive treatments are proven successful. The range of brain areas available for study in these participants is diverse since electrodes are placed within a variety of cortical (e.g., orbitofrontal, motor, temporal cortices) and/or subcortical (e.g., medial temporal lobe, basal ganglia) areas depending on an individual’s clinical prognosis. Patient populations with implanted neural devices thus provide a rare scientific opportunity to directly record from and stimulate a variety of locations within the human brain during freely moving behaviors without the confounds of immobility and motion-related artifacts present in other recording methods [7-9]. A few neuroscientists have begun to capitalize on this opportunity [10][11][12]. However, these recent studies did not provide universally applicable methods for real-time viewing and control nor the ability to stimulate and perform precise synchronization of the intracranial electroencephalographic (iEEG) and externally acquired data during free movement. Here, we provide a first-of-its-kind mobile deep brain recording and stimulation (Mo-DBRS) platform that enables flexible wireless control over chronically implanted neural devices and synchronization with wearable technologies that can record heart rate, respiration, skin conductance, scalp EEG, full-body movement/positional information, and eye movements. Moreover, when used in conjunction with virtual/augmented reality (VR/AR) technologies, the Mo-DBRS platform can provide ecologically valid environments to simulate real-world experiences and open a new area of research in the fields of basic, cognitive, and clinical neurosciences.

We implemented, characterized, and validated the components of our Mo-DBRS platform in five research participants (Table S1) previously implanted with the RNS® System (NeuroPace, Inc., Mountain View, Fig. 1a) with electrodes in a variety of medial temporal and frontal regions (Fig. 1b, Table S2) for treatment in accordance with the product labeling. All participants volunteered for the study by providing informed consent according to a protocol approved by the UCLA Medical Institutional Review Board (IRB). A central part of the platform is a set of Research Tools (see online methods for details), which allows for wireless and programmable control of the RNS System, including the implanted neurostimulator. The Mo-DBRS components can be carried by backpack in a full (weight = ∼ 9 lbs, Fig. 1c, d, S1a) or lightweight (Mo-DBRS Lite; weight = ∼ 1 lbs, Fig. S1b, Fig. S2) wearable platform for use during ambulatory (Fig. S1a) or stationary (Fig. S1c) research paradigms.

**Fig. 1.**
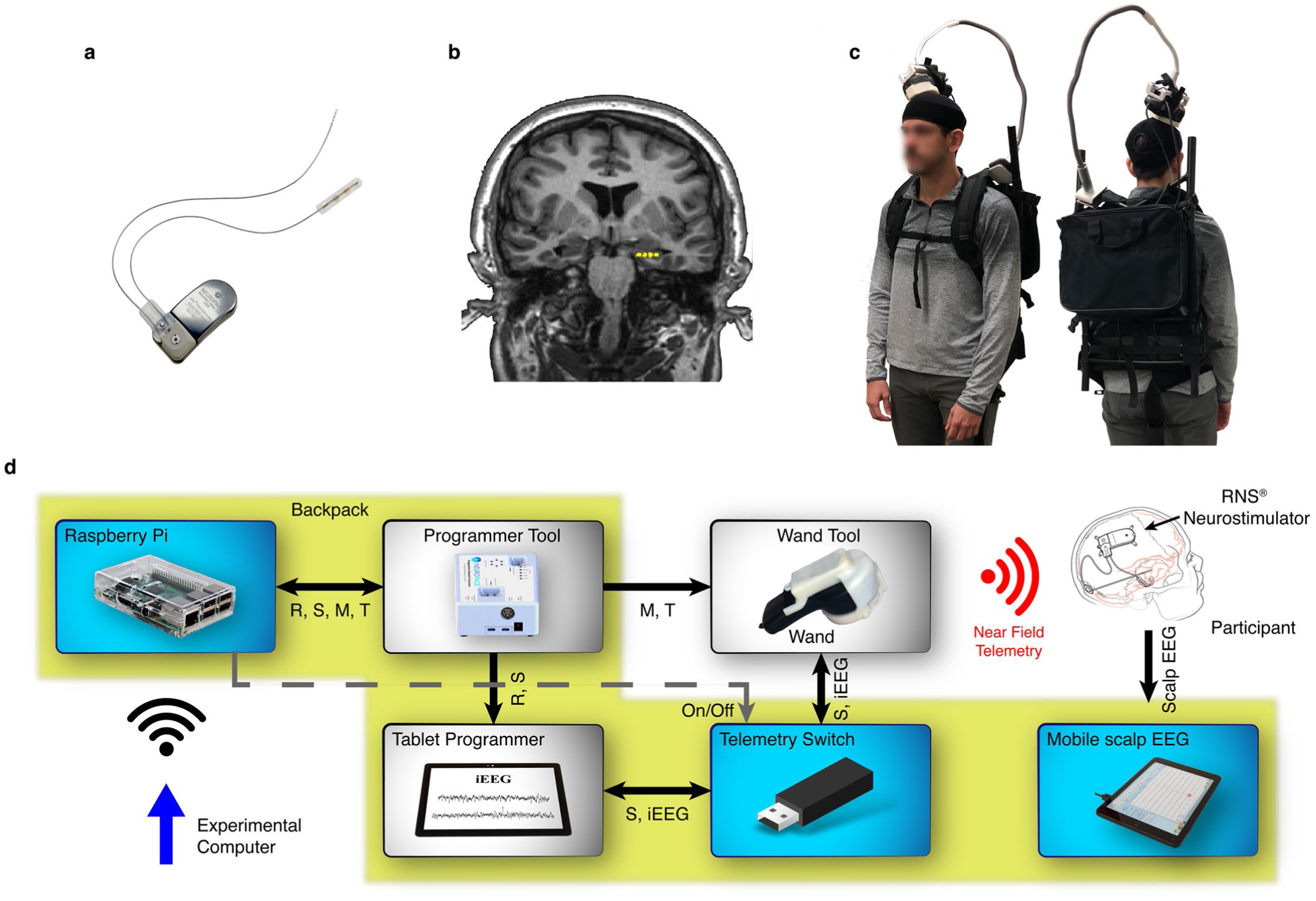
Mobile Deep Brain Recording and Stimulation (Mo-DBRS) platform. **a**, Example RNS Neurostimulator model RNS-300M with two leads containing a total of eight depth electrode contacts. **b**, An example participant’s magnetic resonance imaging (MRI) scan showing four-electrode contact locations (yellow) in the left hippocampus (electrode locations determined by co-registering a post-implant computerized tomography image (CT). **c**, Full platform with wearable backpack that carries the **d**, Mo-DBRS Research Tools, which receive commands from an Experimental Computer, including R – *Store*, S – *Stim*, M – *Magnet*, T – *Mark*. A Raspberry Pi serves as an input to the Research Tools, via wireless forwarding of commands to the Programmer Tool. The Wand wirelessly (Near Field Telemetry) transmits the commands and receives data to and from the implanted RNS System. Items highlighted with a yellow background are Research Tools that can be carried in a wearable backpack. Shown is the Tablet Programmer; however, the same setup applies if using the Laptop Programmer. Solid arrows indicate a USB serial connection. A dashed arrow indicates a single wired connection. The black or red Wifi symbols indicate a wireless local network connection or Near Field Telemetry, respectively.

The full wearable version of the platform (Mo-DBRS) provides real-time viewing, storage, and synchronization of iEEG with external data, as well as on-demand triggering of deep brain stimulation (DBS). These functionalities can be accessible on a local wireless network through a TCP/IP Socket server running on a small board computer, Raspberry Pi (RP). The implanted RNS Neurostimulator communicates via Near Field Telemetry, thus requiring the Wand to be placed on a participant’s head, close to the underlying implanted RNS Neurostimulator (Fig. 1c). The Programmer, when used with the custom-built Programmer Tool, can accept commands for iEEG storage (Store command) and DBS (Stim command). The Wand, when used with the custom-built Wand Tool, allows for the injection of a signal (Mark command) into the Real-Time iEEG data that can be used for synchronization. The Wand Tool can also deliver an electromagnetic pulse (via a Magnet command), which triggers the local storage of iEEG on the implanted RNS Neurostimulator itself. While the Real-Time iEEG data can be transferred to the Programmer in real-time via the Store command, the Magnet command triggers the iEEG data to be saved on the implanted RNS Neurostimulator and then be retrieved at a later time via the Wand. For remote programmable control of the RNS System, the RP can send commands to the Programmer Tool, which are forwarded to the Programmer (Real-Time iEEG Store and Stim) and the Wand Tool (Mark and Magnet). The use of the RP timestamp logs and the Mark command allows for the synchronized time difference between the RP’s input and the Real-Time iEEG data to be less than 16 milliseconds (Fig. S3, Fig. S4). During experimental task paradigms, the Real-Time iEEG can be remotely viewed from the Programmer, via screen-sharing programs over the network (Figs. S5b, S6).

The lightweight version of the platform (Mo-DBRS Lite) uses onboard storage capabilities of the implanted RNS Neurostimulator and the ability to trigger a Magnet command with a single electromagnet device (Fig. S2a), resulting in a recording-only solution. Mo-DBRS Lite thus requires a separate stand-alone electromagnet device with sufficient driving power that can send a Magnet command, which can be triggered 1) with a physical button press, 2) at a pre-configured repeated number of programmable seconds (Fig. S2a), 3) or be fully programmable (Fig. S2b) via a RP server that sends Magnet commands on-demand once a selected message is received from the local network. For all three options, a visible LED worn externally (Fig. S2b) is triggered simultaneously with the Magnet and can be used to synchronize the stored iEEG activity offline with data acquired from external wearable devices. For the 3rd option, the RP timestamp logs can be used in addition to the Magnet command and LED for synchronization purposes.

One unique aspect of the Mo-DBRS platform is that it can be used with human participants who can wear on-body sensors and devices (wearables). We, therefore, include in the platform, solutions for synchronization of iEEG, DBS, and wearable technologies including full-body motion capture (Figs. 2b, S1e-f), eye tracking (Figs. 2c, S1a,c), biometrics (heart rate, skin conductance, respiration, Fig. 2d, S1d), and scalp EEG (Fig. 2f). In addition to traditional 2-D computer-based tasks (Fig. S1c) and real-world scenarios (Fig. S1a), the Mo-DBRS platform can also allow participants to view synchronized task stimuli on wearable VR/AR headsets (Figs. 2a,h, S1e-f) that simulate naturalistic real-world experiences, therefore, allowing for full head and body movements under experimental control.

**Fig. 2.**
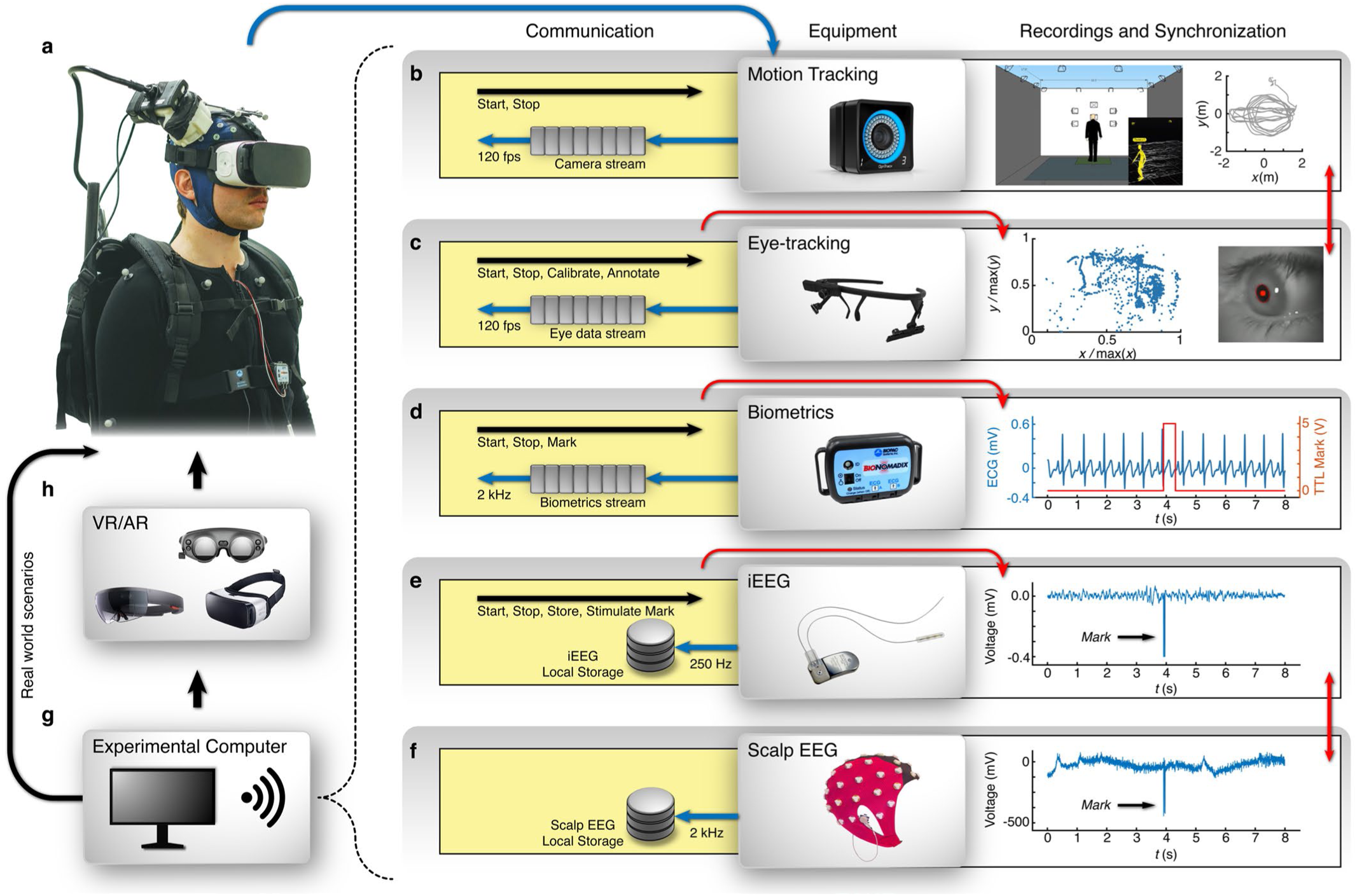
Wearable technologies included in the Mo-DBRS platform. On the right (**b** – **f**), the wearable measurements are listed, including their wireless connection with the Experimental Computer, a synchronization solution, and an example data trace for each. Yellow boxes reflect the communication flow between the Experimental Computer and the external wearables. Within the yellow boxes, the black arrows show the communication flow (commands sent), blue arrows show the data flow (gray boxes indicate real-time data transfer and gray circles indicate local on-device storage), and red arrows point to one possible synchronization solution between different components. **a**, An example participant wearing the full Mo-DBRS platform with a VR headset (also works with an AR or eye-tracking headset). **b**, Wall-mounted cameras for 3-D full-body motion tracking used to record position and movement streaming in real-time at a rate of 120 fps. Cameras are connected through the local wireless network, and a preconfigured recording is controlled with a start/stop function. In the recording box, there is an example real-time view of the participant’s location (left) and an extracted top-down view of the participant’s movement (right). **c**, Wearable eye-tracking system with 120 fps head direction camera and 200 fps eye cameras used for real-world environment studies. Eye-tracking is synchronized with wearable or wall-mounted motion capture cameras from within the Experimental Task Paradigm controlled via the Experimental Computer. The eye-tracking headset is connected through the local wireless network awaiting start, stop, calibrate, or annotate commands. In the recording box, there is a snapshot view from one eye tracking camera (right), and a 2D projection of pupil movement (left). **d**, The wearable biometric system for heart rate, skin conductance, and respiration measurements. In the recording box, there is an ECG recording trace in line with a synchronization signal containing a 400 ms wide synchronization pulse. **e**, The wireless implantable RNS Neurostimulator, which is connected/synchronized with the Experimental Task Paradigm controlled by the Experimental Computer through the local wireless network via the RP. In the recording box, there is an example of a raw Real-Time iEEG data trace with an example *Mark* command signal used for synchronization. **f**, Wearable scalp EEG cap. In the recording box, is an example 2000 Hz sampled scalp EEG data trace with an example *Mark* command signal used for synchronization. **g**, Experimental Computer (e.g., laptop, tablet, or phone) running the Experimental Task Paradigm. **h**, VR/AR headsets integrated and synchronized that are shown include the SMI Samsung Gear VR, Microsoft HoloLens, and Magic Leap, but others can also be used (HTC Vive, Oculus, etc). For studies done in a real-world environment, **h** step can be bypassed, and eye-tracking goggles shown in **c** can be used instead.

Currently, researchers use a variety of stimulus presentation software programs (e.g., Matlab, Python, Unity, etc.) on an Experimental Computer (Fig. 2g) to implement their Experimental Task Paradigms whether stationary or mobile. Nearly all of the software programs allow for script modifications where a TCP/IP Socket client can be added to establish a real-time connection and control over an RP. Thus, the Mo-DBRS platform includes open-source code solutions for the most commonly used programming environments (i.e., Matlab, Unity, and Python), allowing experimenters to adapt and upgrade their Experimental Task Paradigms to work with the Mo-DBRS platform. We also provide a graphical user interface solution for use on a phone/tablet (Fig. S5) in the case of manual non-automated tasks. It must be borne in mind that the specific wearable platform version that is utilized (i.e., Mo-DBRS or Mo-DBRS Lite) affects the system setup and, as such, the tradeoffs must be considered during the design of the Experimental Task Paradigm (see Supplementary Information and Table S3).

All wearable equipment is connected to the same local network, and is synchronized with the Experimental Task Paradigm, by using software timestamping, *Mark* commands, and other task-dependent visual or audio events captured by wearable recording devices. The total synchronization accuracy for the Mo-DBRS platform is ∼ 16 ms. An example participant wearing the full Mo-DBRS platform is shown in Figure 2, with the underlying code structure in Fig. S13. However, most Experimental Task Paradigms seldom require all of these components simultaneously and, therefore, can be customized for a specific research study (Fig. S1).

While iEEG allows for recording of activity within specific deep brain structures, scalp EEG remains a prominent method to probe the human brain, as it is a more readily available methodology due to its non-invasive nature [9][13]. The Mo-DBRS platform allows for simultaneous iEEG and scalp EEG (Fig. S7)—an opportunity that can elucidate a link between deep and surface brain activity and bridge findings across studies. To enable simultaneous iEEG and scalp EEG, we developed two solutions for minimizing RNS System telemetry-related artifacts in scalp EEG data through the use of a programmable switch for disabling/enabling telemetry (Telemetry Switch, Figs. S8, S9) and signal processing methods for noise reduction (Online Methods, Figs. S10, S11, S12).

Currently, in the fields of Cognitive and Clinical Neuroscience, naturalistic behavioral paradigms with rich behavioral data limit the recording of neural data, while paradigms comparatively rich in neural data are almost impossible to carry out in naturalistic settings. The proposed platform provides a new method for modulating and interrogating the human brain during naturalistic behaviors with ecologically valid tasks by enabling wireless and programmable DBS and iEEG recordings synchronized with biological and behavioral data via wearable technologies. While there are clear advantages in adopting the platform in order to provide a unique window into the human brain, new users should be cognizant of limitations related to iEEG recordings in patients with neurologic or psychiatric disorders [9]. We provide the functionalities and details necessary for building and optimizing the Mo-DBRS platform— during ambulatory or stationary behavioral paradigms—through real-time wireless control of sensing, stimulation, and synchronization with external devices such as eye-tracking/VR/AR headsets and other external behavioral measurements. Several components of the platform including open-source code provided for wireless programmable synchronization with wearables sensors can be adapted for use with other existing neuroprosthetic devices such as the Medtronic RC+S or others in development. In addition to enabling basic Neuroscience studies, when used in patients with neurologic and psychiatric disorders along with continuous behavioral metrics, the Mo-DBRS platform provides an opportunity to characterize neural mechanisms and develop and test novel treatments, in unprecedentedly data-rich and naturalistic environments.

## Online content

See online methods for full details required to build the platform, including example code for remote control and synchronization of components (https://github.com/suthanalab/Mo-DBRS).

## Acknowledgements

We thank Nader Shaterian and FabLab (Golden Beach, FL), Interactive Lab (Moscow, Russia), Merit Vick (NeuroPace, Inc.), and Jason Travis (NeuroPace, Inc.) for technical assistance. This work was supported by UCLA startup funds, the National Institute of Neurological Disorders and Stroke (NS103802; NS058280), the McKnight Foundation (Technological Innovations in Neuroscience Award), and the Keck Foundation. We also thank the participants for taking part in our study. N.R.H. is an employee of Neuropace, Inc.

## Online Methods

### Mo-DBRS Platform Components and Setup

We outline below detailed methods needed to re-create the Mo-DBRS platform as well as suggested solutions for proper use, setup, and synchronization of data streams. We also describe the procedures used to validate the platform capabilities as well as characterize synchronization latencies. Corresponding findings are presented in Supplementary Information. Required code and algorithms are publicly available as open-source code and can be downloaded from GitHub (https://github.com/suthanalab/Mo-DBRS).

#### Intracranial EEG (iEEG) Data Acquisition

The FDA approved RNS System (Fig. 1a) includes an implantable neural device (RNS Neurostimulator) used to detect abnormal electrical activity in the brain and respond by delivering imperceptible levels of electrical stimulation to normalize brain activity before an individual experiences seizures. Each participant with an RNS System has one or two implanted depth electrode lead(s) that are 1.27 mm in diameter, each with four platinum-iridium electrode contacts (surface area=7.9 mm^2^, 1.5 mm long) with an electrode spacing of either 3.5- or 10-mm. To localize electrode contacts to specific brain regions, a high-resolution post-implantation CT image is obtained and co-registered to a pre-implantation whole brain and high-resolution MRI for each participant using previous methods [1] (Example Fig. 1b). The RNS System records iEEG activity on up to four bipolar channels at 250 Hz. Onboard Analog filters can be configured to capture the widest possible bandwidth 4 – 120 Hz for the RNS-300M or 1 – 90 Hz for the RNS-320 models. The acquired iEEG data can be stored in real-time (Real-Time iEEG) or transferred at a later time (Magnet-triggered iEEG) to a Laptop or Tablet Programmer device (see *Mo-DBRS Research Tools* below for details). To view Real-Time iEEG activity, a Virtual Network Computing (VNC) TightVNC server was installed on the Laptop Programmer, running while placed in the participant’s backpack. Until VNC can be supported on the Tablet Programmer, a phone camera with TeamViewer installed can be used for viewing the real-time data (Fig. S6).

#### Virtual and Augmented Reality (VR/AR)

The Mo-DBRS platform currently enables successful synchronization with VR/AR headsets equipped with eye-tracking, including the SMI Samsung gear, TOBII HTC Vive, Microsoft HoloLens, Magic Leap and can be adapted for use with other VR/AR headsets. VR/AR environments are programmed using openly available 3-D modeling and game development tools such as the Unity game engine and the C# language, to implement customized immersive environments with controlled stimuli and functionalities. Using motion capture (wearable or wall-mounted cameras), the participant’s location can be mirrored in real-time in the VR/AR application. Example code for this implementation is openly available on GitHub (https://github.com/suthanalab/Mo-DBRS).

#### Biometric Measurements

Simultaneous Photoplethysmogram (PPG), electrodermal activity (EDA), electrocardiogram (ECG), and respiration (RSP) measurements can be performed using the wireless and wearable Smart Center system (BIOPAC® Systems, Inc.), controlled by the Acq*Knowledge* software interface. The Mo-DBRS platform integrates the BioNomadix Smart Center device, a digital interface with USB connection, that collects data from two BioNomadix (Fig. 2d) transmitters (First: RSP + ECG; Second: PPG + EDA). The USB-TTL Interface module can be used for millisecond accurate TTL synchronization through a USB serial port.

The setup uses three ECG recording electrodes fixed to the left and right upper chest and one on the left lower chest. Two EDA recording electrodes can also be placed on the tips of non-dominant hand fingers (Fig. S1d). The BioNomadix and USB-TTL interface module is connected to acquisition modules and an Experimental Computer. Within the Biopac Acq*Knowledge* software, which is run on the Experimental Computer, all recordings can be configured with a sampling rate of 2 kHz. The Biopac acquisition software can run throughout the experimental session, and the data can be synchronized with the iEEG data and other recordings from the Mo-DBRS platform offline.

#### Eye-tracking

Many VR/AR headsets currently have built-in eye-tracking (e.g., SMI Samsung gear, TOBII HTC Vive, Microsoft Hololens, Magic Leap), which is automatically synchronized with the VR/AR tasks (for example programmed in the Unity game engine). For real-world tasks in naturalistic environments, the lightweight Pupil-Labs eye-tracking device (Pupil Core headset) [2] can be used, which has an open-source platform for pervasive eye-tracking and mobile gaze-based interaction -- providing binocular eye cameras (up to 200 fps), and an external world-view camera (up to 120 fps) (Fig. 2c). An Experimental Computer can control the eye-tracking hardware through ZeroMQ asynchronous messaging over a local network. The Pupil-Labs mobile application and an Android phone that controls the head-mounted eye-tracking device can be used for ambulatory tasks while a direct connection is sufficient for stationary tasks. The Pupil Capture plugin manager can be configured to include: Annotation Capture (software synchronization marks), Blink Detection (online blink detection algorithm), Pupil Remote (allows wireless eye-tracking control and streaming), Time Sync (for network clock synchronization). For calibration, we used the Screen Marker Calibration and Accuracy Visualizer to assess its quality.

#### Scalp EEG

Participants with chronically implanted neural devices can also wear a scalp EEG cap that allows for ambulatory behaviors. We integrated with the Mo-DBRS platform a mobile 64-channel scalp EEG system (Wave Guard and eego™ mylab system, ANT Neuro, The Netherlands) that includes a lightweight amplifier (∼ 2 lbs) which connects to the cap and a small tablet, which can both be carried in a backpack to which data is being transmitted. After the experimental session, electrode digitization can be completed to verify scalp EEG electrode positions relative to the head and underlying brain (if MRI is available). Scalp EEG and iEEG data can then be synchronized (see Mo-DBRS platform synchronization section below) and analyzed offline.

#### Mo-DBRS Research Tools

Flexible control over the implanted RNS Neurostimulator, real-time iEEG recording and storage, deep brain stimulation (DBS) delivery, and synchronization of data streams during free movement, requires additional tools that the user can re-create. These Research Tools are listed below. Previous studies [3][4][5] have used variations of the Programmer, Programmer Tool, Wand, and Electromagnet (see components 2 – 5 and 8 below). Components with a single asterisk come with the clinical RNS System, and those with a double asterisk are Research Tools that were provided by NeuroPace, Inc. The Mo-DBRS platform requires components 1 – 7. The Mo-DBRS Lite platform requires component 8, while component 1 is optional.

1. **Experimental Computer** An experimental device (e.g., laptop or phone) that runs the behavioral or cognitive task of interest (Experimental Task Program) and sends commands over the network to remotely control the RNS System (Fig. 1d).
2. **Programmer\*** A Programmer (Laptop or Tablet version) that comes with the clinical RNS System can bewas used to retrieve, store, and monitor Real-Time iEEG as well as trigger delivery of DBS (Fig. 1d). The Laptop Programmer is only compatible with the older RNS-300M model, and the Tablet Programmer is compatible with both the older RNS-300M and newer RNS-320 models. The Tablet Programmer does not require the researcher to manually program data storage commands (i.e., the *Store* command, see *Programmer Tool* section below) into their Experimental Task Program (see GitHub link for example code), unlike the Laptop Programmer, since Real-Time iEEG data storage on the Tablet is performed automatically in chunks of a predefined duration (programmable by user) between 60 seconds and 90 minutes. Lastly, the Tablet Programmer is more responsive and thus delivers commands to the implanted RNS Neurostimulator faster (see *Mo-DBRS Platform characterization and validation* section) compared to the Laptop Programmer.
3. **Programmer Tool\*\*** To control the Programmer’s graphical user interface, an Arduino SAM Board (Model “Due”, 32-Bit ARM Cortex-M3) can be used, which accepts one ASCII byte from the Experimental Computer and forwards the information to the Programmer or Wand Tool (Fig. 1d). There are two firmware versions for the Programmer Tool, one compatible with the Laptop Programmer and the other one with the Tablet Programmer. The Laptop Programmer Tool allows for a trigger of four valid commands on the Programmer: 1) The *Store* command stops Real-Time iEEG transmission, stores up to 240 seconds of previously observed data, and then starts Real-Time iEEG again via a mouse click that selects related functions within the Laptop Programmer graphical user interface. 2) The *Stim* command initiates the Laptop Programmer to trigger delivery of DBS under predefined parameters that can be manually set by the user on the Laptop Programmer. 3) The *Mark* command delivers a timed and visible pattern into the Real-Time iEEG recording, allowing for synchronization with externally acquired data. Specifically, the *Mark* command triggers the Wand Tool to inject a distinctive noise pattern, 64 ms in duration, into the Real-Time iEEG data (Fig. S3a) that can be analytically distinguished from the ongoing neural signal (see *Mo-DBRS platform synchronization* section below). 4) The *Magnet* command delivers a 520 ms wide electromagnetic pulse (Fig. S3b), which triggers iEEG storage on the implanted RNS Neurostimulator. This command allows for an alternative way of storing the iEEG data through the Wand Tool that does not require real-time transmission to the Programmer. However, if using the *Magnet* command to trigger data storage, the iEEG data needs to be externally downloaded (by interrogating the implanted RNS Neurostimulator via the Wand) before it gets overwritten by another *Magnet* command (given the RNS System’s limited storage space). The RNS-300M (320) model can store up to 7.5 (13) minutes of Magnet-triggered iEEG data (from 8 electrodes, 4 bipolar recordings). The RNS System data buffer capacity is increased if recording on fewer channels (e.g., up to 30 minutes [RNS-300M] and 53-minutes [RNS-320] recordings on 1 bipolar channel). The Tablet Programmer Tool provides a broader range of possible controls in addition to the described four commands enabled with the Laptop Programmer. Specifically, there is added support for: 300M/320 mode selection, independent Real-Time iEEG start/stop commands, and software labeling for synchronization and other purposes. The Programmer Tool’s input is a USB serial connection (baud rate: 9600 bps Laptop Programmer Tool, 57600 bps Tablet Programmer Tool). It has two outputs, an output USB connection towards the Programmer and a proprietary NeuroPace connection towards the Wand Tool (described below, Fig. 1d). The Programmer Tool is powered with the input USB connection or with a 12 V battery (required for *Magnet* commands).
4. **Wand\*** A device that comes with the clinical RNS System that every patient has in their possession and is used to communicate wirelessly with the implanted RNS Neurostimulator via Near Field Telemetry when placed on the surface of the head (Fig. 1d).
5. **Wand Tool\*\*** A device that holds the Wand and produces the command triggered *Mark* pulses for Real-Time iEEG synchronization and *Magnet* pulses for Magnet-triggered iEEG storage (Fig. 1d).
6. **Wireless Control Device (Raspberry Pi)** All of the previously described Research Tools can be used for stationary (tethered) laboratory computer-based Experimental Task Programs. However, to enable a completely wearable solution that allows for free movement such as during spatial navigation of VR/AR or real-world environments, a small single-board computer, such as the Raspberry Pi 3 (RP, Fig. 1d), Model B, running Raspbian GNU/Linux 9.9 (Stretch) distribution, can be used as a wireless bridge between the Experimental Computer and Programmer Tool. The RP was chosen because it satisfies the minimal requirements: onboard wireless (e.g., network controllers, Bluetooth, or others) and USB peripherals. Basic RP functionality involves running a secure TCP server that forwards commands from the Experimental Computer to the Programmer Tool. If tasks are completed in indoor environments, the RP can be configured with a static IP address and connected to a local wireless network. For our experimental setup, we have used the Asus RT-AC5300 router; we suggest using this router or one with similar performance. For tasks in outdoor environments, the RP can be configured to provide a remote Access Point (Wi-Fi hotspot) [6]. If not explicitly stated otherwise, all example scripts (Github) on the RP are run using Python 2.7 and are available for download. The scripts on the RP run the TCP server to which clients from the Experimental Computer can be connected. Once the connection is established, the server can be put into an idle state (i.e., blocking the read call function) until a command from the client is received. When the acknowledge receipt is received back from the Programmer Tool, a timestamp is logged. Experimental Computer timestamps are also logged before each command, which can be later used to verify synchronization methods. The role of the RP is critical in the following cases: 1) During the Experimental Task Program, the RP is the only communication channel between the Experimental Computer and the Programmer Tool–this connection is necessary to send behaviorally relevant commands to the Programmer Tool; 2) When the Experimental Task Program requires complex command sequences and their delivery at specific times with high precision. In general, partial implementation of these functions at the RP level, rather than on the Experimental Computer, warrants better flexibility and accuracy of timing; 3) When the Experimental Task Program can be entirely implemented on the RP and, thus, the RP can serve the role of the Experimental Computer.
7. **Telemetry Switch** Since the RNS Neurostimulator is a chronically implanted neural device with no externalized wires, it is possible to simultaneously record synchronized scalp EEG (see *Mo-DBRS platform synchronization* section for details). We provide solutions for noise reduction in the simultaneously recorded scalp EEG. The noise is a result of the Wand’s telemetry given that the scalp EEG cap has to be placed in between the Wand and the implanted RNS System. The first solution is the use of the *Magnet* command to trigger iEEG storage, which does not require telemetry to be implemented and thus results in artifact-free scalp EEG recordings. However, this method lacks the temporal precision required for some Experimental Task Programs since the latency is greater than 250 ms. Therefore, the second solution that can be used is to programmatically switch on/off telemetry (experimental Telemetry Switch, via a USB connection) to prevent telemetry artifacts when possible. While this functionality is possible by manually unplugging the Wand USB cable from the programmer, we instead built a custom Telemetry Switch that can be placed between the Programmer and the Wand (Fig. 1d) to enable/disable telemetry when needed (i.e., enabled when sending commands) by controlling the digital input connected to the RP. If scalp EEG is not needed, the switch may remain ON. For circuit details and code required to build the Telemetry Switch, see Fig. S8 and GitHub. Telemetry Switch usage is described in the section on *Scalp EEG Artifact Reduction – Telemetry Switch*. An additional solution is to use artifact rejection offline after data acquisition (see section below on *Scalp EEG Artifact Reduction – Offline Processing*).
8. **Electromagnet\*\*** The Mo-DBRS Lite version of the platform can be used as an alternative solution to the full Mo-DBRS platform, through the use of the custom-built experimental electromagnet device, which, due to its size, is a more lightweight solution and thus comfortable for participants who may be physically impaired and/or have limited mobility. The Mo-DBRS Lite platform requires an electromagnet device that produces a pulse with minimum duration of 520 ms (anything shorter cannot be detected by the implanted RNS Neurostimulator) that can be triggered by the custom-built battery-powered wearable control box (see Fig. S2 for circuit details required to reproduce the control box). The electromagnet component (Fig. S2b-c) is a Research Tool provided by NeuroPace. Depending on a predefined length of Magnet-triggered stored iEEG data specified by the user in the Programmer prior to the experimental session, the electromagnet device can be set to be triggered at predefined configurable time intervals (e.g., 30/60/90/180 or 240 s in our case). A small LED—located at proximity of the electromagnet—can be configured to turn on simultaneously with the triggered electromagnetic pulse, and thus be captured by external or wearable cameras, which can later be used to synchronize iEEG activity with external data. Manual triggering of the electromagnet and timers can be handled using a PIC controller (see assembly code on GitHub) that requires additional circuitry for power amplification to drive the electromagnet (Fig. S2a,b). For remote and programmatic control of the wearable electromagnet device (including pulse duration and time of delivery), additional circuitry can be added to the RP (see Fig. S2a,c for full details needed to reproduce).

### Mo-DBRS Platform characterization and validation

We tested the Mo-DBRS platform in-vivo in five participants previously implanted with the RNS Neurostimulator (Table S1) for treatment in accordance with the product labeling and ex-vivo (benchside) with a test RNS Neurostimulator (see Supplementary Information for results). For in-vivo tests, all participants volunteered for the study by providing informed consent according to a protocol approved by the UCLA Medical Institutional Review Board (IRB). To avoid unintended stimulation artifacts in the iEEG activity, stimulation was turned off with the participant’s consent and the RNS System was used only to record brain activity during experimental sessions.

#### In-vivo testing

Five participants wore the Mo-DBRS platform and maneuvered freely through an indoor environment, where wall-mounted motion capture cameras were used to monitor position and full-body movements with sub-millimeter precision (Fig. 2b). Eye position and movements were recorded with the Pupil Core eye-tracking headset (Fig. 2c) worn and carried with the Pupil Mobile phone device inside of a wearable backpack (Fig. 1c, Fig. S1a). Heart rate (ECG) was measured from the participant’s chest, and skin conductance via sensors from the fingers (Fig. 2a,d, Fig. S1d). Participants also wore a scalp EEG cap and were asked to carry a sturdy backpack in which we placed necessary equipment (e.g., Mo-DBRS Research Tools, scalp EEG amplifier and data acquisition tablet). Participants were able to wear the setup comfortably for several hours throughout the day. The Wand and Wand Tool were tightly strapped together with a ‘Wand holder’ (iPhone holder was used; Fig. 1c, Fig. 2a, Fig. S1) using Velcro tape and a rubber band to ensure a stable connection and prevent misalignment between the Wand and Wand Tool, which if not done properly would have led to missed *Marks*. The flexible metal Wand holder was secured to the backpack to relieve the weight of the Wand and Wand Tool on the participant’s head and to provide stability and movement flexibility. Lastly, the Wand was angled close to the implanted RNS Neurostimulator location, and once Real-Time iEEG telemetry was established, it was fixated to the scalp EEG cap using Velcro tape placed on the side of the Wand Tool that was secured to the participant’s head (Fig. 1c, Fig. 2a, Fig. S1). We tested both the full Mo-DBRS and Mo-DBRS Lite versions of the platform, which differed in how iEEG data was handled (see Table S3 and *Mo-DBRS (Real-Time iEEG) versus Mo-DBRS Lite (Magnet-triggered iEEG) trade-offs* section in Supplementary Information for results). Mo-DBRS uses the *Store* command to save streamed Real-Time iEEG on the Programmer (Fig. 2e) and the *Mark* command for synchronization. The Mo-DBRS Lite version, on the other hand, uses the *Magnet* command to save iEEG on the implanted RNS Neurostimulator and the same *Magnet* command for offline synchronization. In parallel, the Experimental Computer (Fig. 2g) acquires other streams of data by running the Experimental Task Program (e.g., in Matlab or Unity), eye-tracking software, motion capture software, and biometric measurements (Fig. 2b – d). The Experimental Task Program controls the flow of the experiment based on collected streams of data in real-time. Closing the loop toward the participant in order to control a VR/AR environmental scene, depending on the body and head position, can be achieved by updating the VR/AR headsets (Fig. 2h) or sending Audio/Video messages in real-time. Equipment can be connected through a local wireless network, which allows real-time control, storage, streaming (Fig. 2b-f – communication on the left), and synchronization (Fig. 2b-f – recordings and synchronization on the right) with accuracy of below 16 ms for the full Mo-DBRS platform version. The synchronization fixation points are *Mark* events for the full Mo-DBRS platform and *Magnet* events when using the Mo-DBRS Lite version.

All commands were sent from the Experimental Computer using Unity software (see GitHub for example code). After the experimental session, all data is then collected, synchronized, and scalp EEG can be cleaned using the methods described in the *scalp EEG Telemetry Artifact Rejection* section. In order to characterize the reliability of command delivery, we tested the four wireless commands (*Store, Stim, Mark*, and *Magnet*) using the Research Tools and in the following order:

▪ *Mark* and *Store*: 2 × 240 s iEEG blocks (via *Store* command) with 100 *Mark* commands each, delivered every 2 s.
▪ *Store, Magnet*, and *Stim*: 3 × 240 s iEEG blocks (via *Store* and *Magnet* commands) each with 16 DBS pulses (*Stim* command) with 100 ms burst duration, 0.5 mA output current, each at every 1.8 s, 2 s, and 2.2 s.
▪ *Store, Magnet*, and *Stim*: 3 × 240 s iEEG blocks (via *Store* and *Magnet* commands) each with 16 DBS pulses (*Stim* command) with 2000 ms burst duration, 0.5 mA output current, each at every 3.8 s, 4.2 s, and 5 s.
▪ *Store, Magnet*, and *Stim* with the Telemetry Switch: 2 × 240 s iEEG blocks (via *Store* and *Magnet* commands) each with 8 DBS pulses (*Stim* command) with 100 ms burst duration, 0.5 mA output current, done with the Telemetry Switch on/off, *t*_0_ = 5 s; *t*_1_ = 7 s; *t*_2_ = 3 s; *t*_3_ = 5 s (for *t*_i_ definitions see the *Scalp EEG Artifact Reduction – Telemetry Switch* section below).
▪ *Store, Magnet*, and *Stim* with telemetry switch: 1 × 240 s iEEG block (via *Store* and *Magnet* commands) each with 16 DBS stimulation trials with 100 ms burst duration, 0.5 mA output current, done with the Telemetry Switch on/off, *t*_0_ = 4.8 s; *t*_1_ = 3.9 s; *t*_2_ = 0 s; *t*_3_ = 5 s.

#### Ex-vivo testing

Latency measurements were performed on a test RNS-300M Neurostimulator (bench-side ex-vivo). The same set of commands was sent as in the in-vivo section, but in this case from the RP instead of the Experimental Computer. Command delivery triggered from the RP was estimated by executing a send function while simultaneously changing the state of the RP’s GPIO. The command delivery could be directly observed on the test RNS-300M device’s recording contacts and compared with the RP’s GPIO output using an oscilloscope. The network latencies between the Experimental Computer and the RP were measured directly in code using time libraries (i.e., Python and C language). The Asus RT-AC5300 Wireless Router was used, which provided a reliable connection within the range of 20 meters. Due to different clocks on the test RNS System and external equipment, the RP timestamps and RNS timestamps were synchronized by aligning the first *Mark* appearance in both recordings and setting the RP 1/f slope to the linear fit of the RNS slope.

In addition to characterizing the latencies of individual commands, we also estimated the temporal offsets between the *Mark* command events and the Real-Time iEEG data (i.e., the synchronization/storage accuracy). By sending a simultaneous *Mark* and a voltage pulse (offset at the source = 22.5 ± 41.17 µs) to one of the test RNS Neurostimulator electrode contacts, we were able to measure their relative distance in time (Fig. S3e,f). The voltage pulse was generated on the RP’s GPIO, attenuated from 3.6 V to 2.6 mV using a resistor divider, recorded by analog frontends, filtered, and digitized into the Real-Time iEEG. The Real-Time iEEG also contained *Mark* command artifacts. Ten trials of the *Mark* command showed a discrepancy in the range of 12 – 16 ms (Fig. S3f). This offset came from the recording frontend, digitization (250 Hz sampling), and wireless telemetry.

### Mo-DBRS platform synchronization

We detail here an example solution for utilizing the *Mark* command for synchronization. Specifically, *Marks* are delivered right after the *Store* command, while the Real-Time iEEG data is viewed in real-time to detect any loss of telemetry (Wand connection with the implanted RNS Neurostimulator) in which case the corresponding Real-Time iEEG data can be discarded. *Marks* are then detected using cross-correlation (no normalization) between iEEG data and each of 4 *Mark* signal templates, including a 3-spike template (Fig. S3a) and three versions of a 2-spike template (if 1 of the 3 spikes are missing). *Marks* are identified as the time points where the correlation coefficient between the iEEG data and at least one of the four *Mark* templates is higher than 90% of the maximum determined correlation coefficient. These time points are then used to verify that the corresponding iEEG signal value is at a minimum (maximum absolute value of a signed 10-bit iEEG sample is 512) at the same time points. This cross-correlation procedure is repeated for the three versions of the 2-spike *Mark* template in cases where the full 3-spike *Mark* signal was not captured completely. The *Marks* that are detected using these 2-spike templates are appropriately shifted in time to account for the missed spike given that the predicted time between *Mark* template spikes is known. Using this method, we were able to detect all delivered *Mark* signals in the Real-Time iEEG that contain at least 2 spikes. For synchronization of Magnet-triggered iEEG see the *Mo-DBRS (Real-Time iEEG) versus Mo-DBRS Lite (Magnet-triggered iEEG) trade-offs* section.

For synchronization with the eye tracking system, ZeroMQ API provides annotations for eye-tracking synchronization (∼ 10 ms accuracy) with other streams of data. Software annotations can be delivered from the Experimental Task Program (running the behavioral paradigms) to the Pupil-Labs software (Pupil Capture), which runs in parallel on the Experimental Computer. Annotation makers can be sent to the Pupil-Labs software each time the *Mark* command is sent to the implanted RNS System. Additional and redundant synchronization can be achieved by having a small LED placed on the edge of the outward-facing camera on the eye tracking headset that can turn on for a short period of time (50 ms) simultaneously with the *Mark* command. The LED can be connected and controlled by the RP.

Synchronization, monitored by the Experimental Task Program (Fig. 2, Fig. S13), is summarized as follows:

▪ Depending on the task, the iEEG, biometric measurements, and eye-tracking data can be synchronized using simultaneous *Mark* commands sent by the Experimental Task Program.
▪ iEEG/scalp EEG data can be synchronized with a *Mark* (or *Magnet*) command in the Mo-DBRS (Mo-DBRS Lite) platform.
▪ Due to pipeline delays with the eye-tracking software annotations, an LED indicator can be connected to the RP and turned on whenever a *Mark* or *Magnet* command is being received on the RP (Fig. S13). Similarly to the electromagnet device, we used a 50 ms pulse from the RP to turn on an LED that was captured by the motion capture and eye-tracking cameras.

There are two challenges in terms of synchronizing scalp EEG and iEEG data, as well as minimizing artifacts due to wireless communication with the implanted RNS System (see *Telemetry Switch* section above). Synchronization can be done via the *Mark* or *Magnet* command, as the resulting signal patterns detected in the nearby scalp EEG electrodes are, in fact, beneficial and can be used to align the scalp EEG and the iEEG data streams. On the other hand, noise patterns resulting from DBS or Real-Time iEEG transmission can be avoided using the Telemetry Switch or removed offline using signal processing methods (see *Scalp EEG Telemetry Artifact Reduction* section below).

We performed referential recordings from all accessible channels in the scalp EEG, at 2000 Hz. A higher sampling rate was necessary in order to capture the full frequency range of the RNS Wand telemetry signals in order to model the artifacts that are later used in the cleaning procedure. The referential input signal range is up to 1000 mV_pp_, which was again useful for capturing large telemetry artifacts and for preventing amplifier saturation. For more information on the amplifier specification, see [7].

Scalp EEG data was processed and synchronized with iEEG data using Matlab 2018a with the Wavelet, Signal Processing, and DSP System Toolboxes. We first began with the raw, unfiltered scalp EEG data from all 64 channels sampled at 2 kHz, denoted as **R**_C×Nr_ (C – number of channels; N_r_ – number of sample points per task). Each row of **R** was standard score normalized independently.

Next, we located the distinctive noise patterns (“synchronization artifacts”) that were created by either *Marks* (Fig. S3a) or *Magnets* (Fig. S3b), depending on whether the Mo-DBRS or Mo-DBRS Lite was used. To do this, we created *Magnet* **M**_Nm_ (N_m_ = 1040 points at 2 kHz) and *Mark* **T**_Nt_ (N_t_ = 128 points at 2 kHz) templates using scaled absolute values of their respective waveforms (e.g., Fig. S3b, and Fig. S3a). Similarly to the *Mark* detection in Real-Time iEEG, exact positions of the artifacts were extracted using raw cross-correlation with no normalization between each channel’s time series and templates. For positive side coefficients:

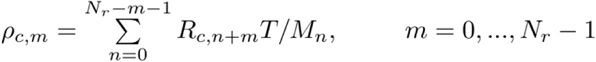

Out of 64 channels, 10 with the highest scaling factor (i.e., standard deviation) were chosen for the synchronization artifact detection due to their proximity to the Wand. Since artifacts across the channels vary only in amplitude, it is best to detect from those with the highest artifact to signal ratio. Additionally, we made three assumptions: 1) each of the 10 selected channels contained all of the delivered *Marks* or *Magnets*, 2) within a single channel the *Marks* and *Magnets* all had the same amplitude, and 3) the scalp EEG signal amplitude during *Magnets* and *Marks* was by far larger than at any other times, including periods of Real-Time iEEG transmission. In practice, *Magnet* scalp EEG artifacts (Fig. S7b) have the largest amplitude, followed by *Mark* scalp EEG artifacts (Fig. S7a). Based on these assumptions, we chose a threshold of 10% lower than the maximum observed correlation per channel. This automated method for synchronization had a 98 % success rate confirmed by using manual inspection of 3 sample scalp EEG/iEEG datasets and comparing to the RP time logs of command delivery as ground truth. The incorrect 2% were all false positives that identified some of the telemetry artifacts as *Marks*. Example figures showing artifacts in scalp EEG used for synchronization can be found in Supplementary Information (Fig. S7).

As each *Magnet* command saves 2/3 of the chosen iEEG data before and 1/3 after the *Magnet* event, we extracted 160 s before and 80 s after the detected *Magnet* timepoint in the scalp EEG data (in this case of preconfigured 240 s Magnet iEEG storage duration). The rest of the data was discarded, and a new scalp EEG matrix **S**_**1**,T×C× Ns_ (T – number of trials; C – number of channels; N_s_ – number of sample points per each 240 s block, e.g., 2000 Hz × 240 s) was synchronized with stored Magnet-triggered iEEG data. When Real-Time iEEG was used, we sent a *Mark* command 1 s before and after the *Store* command. Scalp EEG data in between two detected *Marks*, ∼238 s apart, was extracted as one trial dataset and turned into the same scalp EEG matrix **S**_**1**,T×C×Ns_ (N_s_ –2000 Hz × 238 s), aligned with the Real-Time iEEG data. Lastly, resulting matrices were again standard score normalized per channel. If the *Stim* command was used, iEEG data (stored via *Store* or *Magnet*), could also be synchronized by manual detection of DBS artifacts in iEEG and scalp EEG. Of course, DBS synchronization is only feasible when DBS artifacts are present in the scalp EEG data (Fig S9c, Fig. S7c).

### Scalp EEG Artifact Reduction – Telemetry Switch

We tested the functionality of the Telemetry Switch, which only enabled telemetry communication at specific time points, for instance, when sending a *Store, Stim*, or *Mark* command (*Magnet command* does not need telemetry). Here, despite losing continuous Real-Time iEEG, a *Magnet* command can be used to store the iEEG data since telemetry is disabled. As an example, we performed an experimental session that involved sending a train of DBS bursts, which required cycles of On/Off telemetry. One DBS cycle included enabling telemetry, which took *t*_0_ seconds to get recognized by the Graphical User Interface on the Programmer (scalp EEG remains unaffected by telemetry until *t*_0_). Once telemetry was enabled, the *Stim* command was sent, which required *t*_1_ for delivery. Telemetry was then disabled for an arbitrary period of *t*_2_ immediately after the DBS was delivered, providing clean scalp EEG. The last relevant timing parameter was the time between the DBS cycle preceding the triggering of data storage (*t*_3_) (Fig. S9a). This operation resulted in synchronized iEEG/scalp EEG recordings (Fig. S9b, Fig. S9c). The procedure for sending other commands was done similarly with enabling telemetry (same *t*_0_) and varying *t*_1_ values. Minimal timings were determined in in-vivo experiments (Supplementary Information - *Telemetry Switch characterization and validation* section).

### Scalp EEG Artifact Reduction – Offline Processing

We provide here an offline solution for scalp EEG noise reduction to eliminate any artifacts that remain during enabled telemetry, such as: 1) Telemetry restart artifacts (Type I); 2) DBS artifacts (Type II), and 3) and Real-Time iEEG artifacts (Type III) (Fig. S9c). Type I artifacts appear as a series of alternating segments of two and three spikes, followed by a larger bi-spike (Fig. S10a). Similarly, Type II artifacts consist of segments with two spikes, followed by a bi-spike (Fig. S10b). Spike duration and their relative distance in time were deterministic and fixed– a property that was capitalized upon in our artifact rejection procedure (Spike: 4 samples; bi-spike: 16 samples at 2 kHz). Type III artifacts were spikes at 125 Hz (and harmonics) (Fig. S10c).

Following the automated synchronization method for scalp EEG with iEEG, we normalized the *Magnet*/*Mark*-free scalp EEG **S**_T×C×Ns_. To do this, we first flattened the input matrix to **S**_C×T·Ns_ (in the same order as it came from the raw data in order to simplify analysis) and then applied the same technique to detect Type I and Type II artifacts as we did with *Magnets* and *Marks*. By observing scalp EEG recordings, we constructed binary templates **C**_Nc_ (N_c_ = 3174 points at 2 kHz) and **D**_Nd_ (N_d_ = 2824 points at 2kHz), following respective artifact waveforms from Fig. S9c, 10a, and Fig. S9c, 10b. Note, that 3174 samples or 1.587 s of **C**_Nc_ template plus 2824 samples or 1.412 s of **D**_Nd_ template correspond to a portion of defined *t*_1_ containing artifacts. With no *Marks/Magnets*, we made the same three assumptions used for synchronization and detected two types of artifacts using correlation. Again, for detection, we used ten channels with the highest physical proximity to the Wand. Manual inspection confirmed a 100 % success rate in detection across 3 separate scalp EEG datasets, with RNS time logs of command delivery serving as the ground truth. Once the *Marks/Magnets* were detected, we extracted trials and exact sample points, which resulted in matrices **A**_Na×C×Nc_ and **A**_Na×C×Nd_ (Fig. S10e – 1 and Fig. S10f) for the two types of artifacts (N_a_ corresponds to number of detected artifacts per observed channel). We then applied PCA on scalp EEG N_c_ and N_d_ time series across all channels for each detected artifact separately, while skipping PCA application on adjacent channel pairs [8]. The first 3 PCA components were sufficient to capture the artifacts’ shape shared across different channels. We then eliminated clean scalp EEG during non-spike periods by point multiplication **PC**_i,Nc_ × **C**_Nc_ (and **PC**_i,Nd_ × **D**_Nd_) per each channel and each detected sequence, and applied vertical offset correction for each artifact spike so that it started from 0 (example Fig. S10d). For each segment and channel, we fitted artifact templates to corresponding segments in **A** matrices and subtracted them from it (Fig. S10e – 2). Using previously obtained synchronization timestamps, we reconstructed data into a matrix of original dimensions **S**_**2**,T×C×Ns_. In order to clean Type III artifacts, we filtered each time series within the **S**_**1**_ matrix with a low pass Chebyshev Type I infinite impulse (IIR) filter of order eight and with a cut-off frequency of 125 Hz, and then downsampled data by a factor of 8 from 2 kHz to 250 Hz.

Further artifact rejection can be done using methods reported in [9]. In brief, we used a single channel artifact rejection algorithm in the time-frequency domain. First, the input 250 Hz sampled data were standard score normalized, and then Stationary Wavelet Transform (SWT) was performed (level = 10; Haar base wavelet). Artifact detection was done for approximation and detail coefficients separately. We examined *D*_8_, *D*_9_, *D*_10_, and *A*_10_ coefficients for the input scalp EEG frequency band. Empirically determined thresholds detected outstanding discrepancy from scalp EEG’s approximatively Gaussian distribution for each set of coefficients:

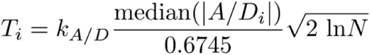

where *N* is number of points in input sequence and *k*_A_ = 0.75, *k*_D_ = 5. Coefficients identified as potentially containing artifacts were thresholded using the Garrote threshold function, after which inverse SWT was applied to reconstruct cleaned signal. This method was applied to each Ns-point time-series within input **S**_**2**,T×C×Ns_, resulting in output **S**_**3**,T×C×Ns_ (Fig. S10e – 3). For more details, see (GitHub) and [9].

Finally, due to high pass filters with low cut-off frequency integrated into scalp EEG equipment, the presence of artifacts caused a voltage drift in raw data (visible slow transients on Fig. S10e – 1,2,3). To account for this we applied IIR high pass filter (order = 8; passband ripple = 0.2; cut-off frequency = 2 Hz) on **S**_**3**,T×C×Ns_ resulting in clean scalp EEG matrix **S**_**4**,T×C×Ns_ (Fig. S10e – 4 and Fig. S10g). To quantify the reduction of artifacts, we calculated the root mean square value (RMS) for **S**_**1**,T×C×Ns_, **S**_**4**,T×C×Ns_, and portions of **S**_**1**,T×C×Ns_ with clean scalp EEG. All RMS values were scaled with a maximum RMS value in **S**_**1**,T×C×Ns_ for given channel (Fig. S10h, i). Additional cleaning results can be seen in Fig. S11 and Fig. S12. Bad channels and channels not containing artifacts were omitted from processing. A total of 17 such channels in the presented case were detected visually and by having SC×T·Ns - CNc/ DNd correlation of less than 0.1 on portions of scalp EEG already identified as artifactual. correlation of less than 0.1 on portions of scalp EEG already identified as artifactual.

