## Supplementary Information for "Wireless Programmable Recording and Stimulation of Deep Brain Activity in Freely Moving Humans"

### Results

#### *Mo-DBRS platform characterization and validation*

We tested the Mo-DBRS platform in-vivo in five participants previously implanted with the RNS Neurostimulator (Table S1) for treatment in accordance with the product labeling and ex-vivo (benchside) with a test RNS Neurostimulator. All participants volunteered for the study by providing informed consent according to a protocol approved by the UCLA Medical Institutional Review Board (IRB). This research demonstrated the functionality of the four commands (*Mark*, *Magnet*, *Store*, and *Stim*) and computed command delivery related latency measurements using distinct Real-Time iEEG artifacts for each command (*Mark* – Fig. S3a; *Magnet* – Fig. S3b; *Stim* – Fig. S3c) and stored the data via the Real-Time iEEG *Store* command (on the Programmer for in-vivo testing). In-vivo we quantified the command latencies during wireless and stationary (tethered) setups. For the wireless setup, the *Mark* and *Magnet* commands were sent from the Experimental Computer to the RP (Fig. 1d) over the local wireless network – TCP/IP protocol. The RP forwarded commands to the Programmer Tool (see online methods for details about Research Tools) over a USB serial connection (baud rate: 9600 bps Laptop Programmer, 57600 bps Tablet Programmer) and then to the Wand Tool over a proprietary custom connection (Fig. 1d). The *Store* command's final point was thus at the Programmer instead of the Wand Tool. The *Stim* command, which had additional connection points (compared to the *Store* command) through the Wand and finally to trigger stimulation within the implanted RNS System (Fig. 1d). Commands that deliver these functions are also characterized by a single character as follows: 1) T – *Mark*; 2) M – *Magnet*; 3) R – *Store*; and 4) S – *Stim*. Total mean latencies from the RP's input to the Wand Tool (sending T, M), Wand (sending S), or Programmer (sending R) output, and thus the execution of the commands, are shown in Fig. S3d. Further, we compared the wireless and tethered setups, where the commands in each case were sent directly to the Programmer Tool over a USB serial connection. Additional delays were introduced by the RP (latency: 2380 ± 142 μs; 1000 trials) and by the network transmission (latency: 149 ± 14 μs; 1000 trials). Lower latencies were measured in the case of the Tablet Programmer Tool, which was due to a higher 57600 bps baud rate and a more responsive Programmer (Fig. S3d). It is worth noting that the commands that involve interactions with the Programmer (*Store* and *Stim*) had much higher latencies due to the lag introduced by the Programmer Graphical User Interface. In practice, a timely execution of these commands required even more conservative timings for proper functionality (Laptop Programmer: *Store* latency=4.6 s, *Stim* latency=2 s; Tablet Programmer: *Store* latency= 2 s, *Stim* latency=1.5 s). These values were obtained by sending multiple consecutive series of the same command (e.g., *Stim* in Fig. S3c).

***Mo-DBRS (Real-Time iEEG) versus Mo-DBRS Lite (Magnet-triggered iEEG) trade-offs***

In general, synchronization is more accurate with the full Mo-DBRS platform that enables Real-Time iEEG using the *Mark* command. The Mo-DBRS Lite platform, which stores iEEG data using the *Magnet* command, does not utilize *Marks* and hence, synchronization depends on extracting the *Magnet* timestamp. As noted prior, a *Magnet* event, once detected in the RNS System, triggers the storage of predefined time period such that 2/3 of the data saved is from before the *Magnet* and 1/3 of the data is saved from after the *Magnet*. The detection of a

*Magnet* is sampled by the RNS System with a 2 Hz frequency and therefore causes a variable offset (up to 0.5 s) between the Magnet triggered and consequently stored iEEG data. To investigate this further, we sent a series of *Marks* and voltage pulse (RP GPIO) pairs (× 5), followed by a *Store* and *Magnet* command in 20 repeated trials (Fig. S4a). The voltage pulse was (Fig. S3e,f) generated from the RP and sent directly to electrode contacts of the test RNS Neurostimulator. *Marks* were used to synchronize recordings with internal timestamps for each event saved on the RP. Voltage pulses were used to synchronize the Real-Time iEEG and Magnet iEEG. The experimental session was controlled from the RP running *C* scripts compiled with *GCC* (version: Raspbian 6.3.0-18). The advantage of *C* compared to Python scripts is that libraries for time measurements, delay, and GPIO control have accuracy and precision of less than 1 μs. Similarly to what was described in the previous section, the first *Mark* was aligned with the first *Mark* timestamp recorded on the RP. Also, the RP time (all *Mark* events) were scaled to match the RNS System clock using linear fitting.

Precision test results are shown in Fig. S4b, c, d. Note that values for *Mark*, *Magnet*, and RP voltage pulse deliveries are significantly lower in this case which is the result of the *C* instruction being executed much faster as compared to Python, thus slowing down the error accumulation that over time can cause events to be picked up by the next iEEG sample (RNS System sampling rate is 250 Hz).

#### *Telemetry Switch characterization and validation*

The experimental telemetry switch was tested and used to partially prevent contamination of scalp EEG with telemetry-associated artifacts. Telemetry restart, DBS, and Real-Time iEEG noise patterns were clearly present during the telemetry enabled state lasting $t_1$ (Fig. S9c, $t_0$ = 5 s;

$t_1$ = 7 s; $t_2$ = 3 s; this sequence allows delivery of continuous DBS every 15 seconds). Due to the inevitable delays in this pipeline, namely recognizing telemetry by the Programmer and stimulation delivery ($t_0 + t_1$), there was a lower limit for how often DBS could be delivered reliably. We determined, empirically, that the minimal timing values were $t_0$ = 4.8 s, $t_1$ = 3.9 s, $t_2$ = 0 s, resulting in a total minimal DBS period of 8.7 s using the Laptop Programmer. Values below this threshold caused either missed *Stim* delivery commands or rejections by the RNS System. The procedure for sending other commands was done similarly with enabling telemetry (same $t_0$) and varying $t_1$ values. It is worth noting that although *Magnet* signals via the *Magnet* command could synchronize scalp EEG with iEEG offline, *Mark* commands and Real-Time iEEG provide much higher accuracy (Fig. S4). Thus, for highly accurate synchronization with scalp EEG, telemetry-enabled blocks of Real-Time iEEG, containing *Marks*, should first be aligned with redundantly stored Magnet iEEG and then aligned with the scalp EEG using either the DBS or *Mark* signals captured in the Real-Time iEEG data. It should also be mentioned that the same method applies to Tablet Programmer, except that the timings are expected to be somewhat lower.

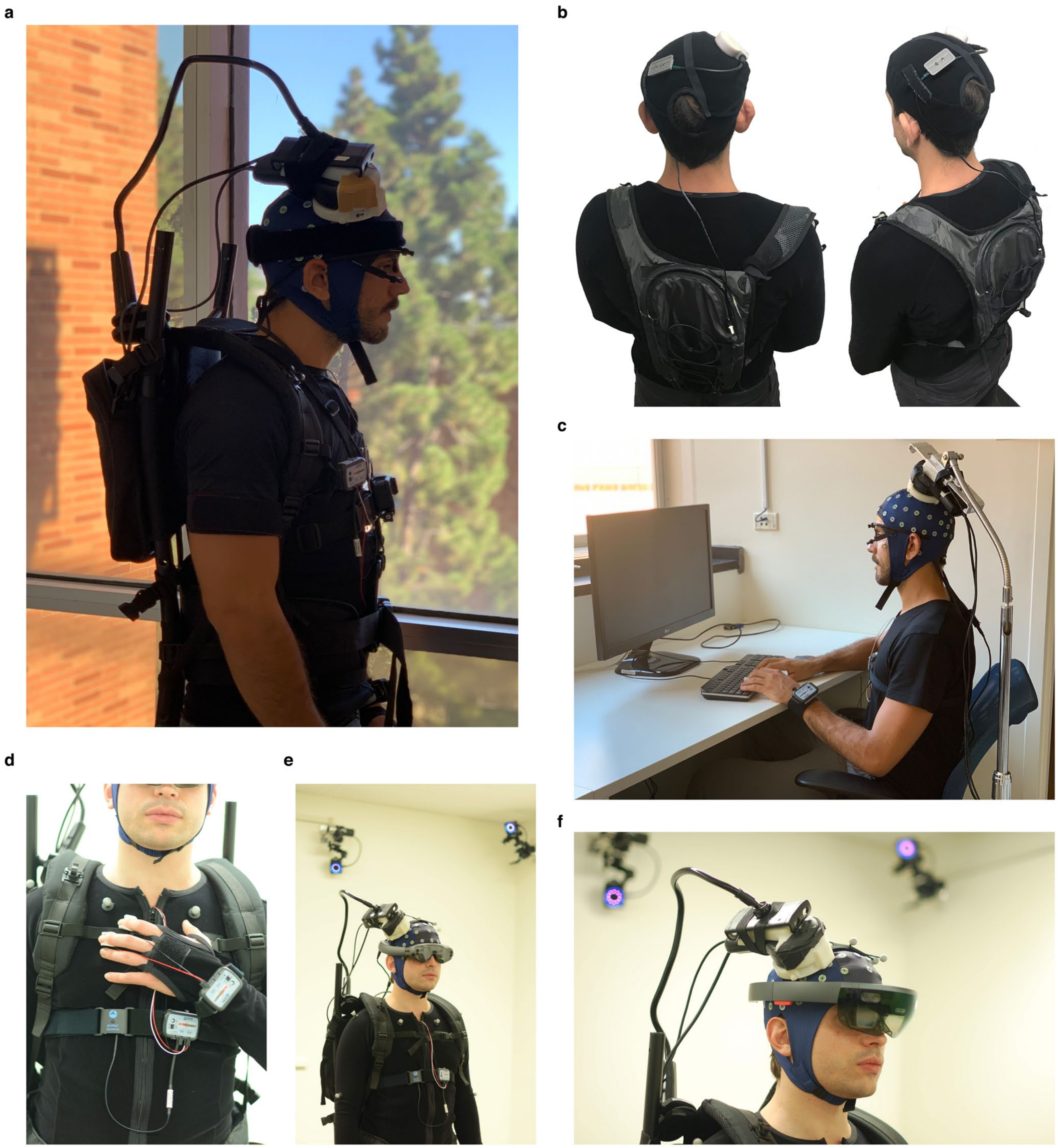

**Fig. S1 | Additional photos of the Mo-DBRS platform in naturalistic and laboratory settings. a,** A participant wearing the Mo-DBRS platform showing the backpack connected to the Wand**. b,** A participant wearing the Mo-DBRS Lite platform showing the backpack with the electromagnet device with head-mounted LEDs used for synchronization. **c,** A participant performing a laboratory-based stationary task. **d**, Close look at the wearable equipment for biometrics including a respiration belt, skin conductance and heart rate sensor. **e**, Participant wearing the Mo-DBRS platform with the Magic Leap AR headset in a laboratory equipped with wall-mounted motion capture cameras. **f**, Same but with the Microsoft Hololens AR headset.

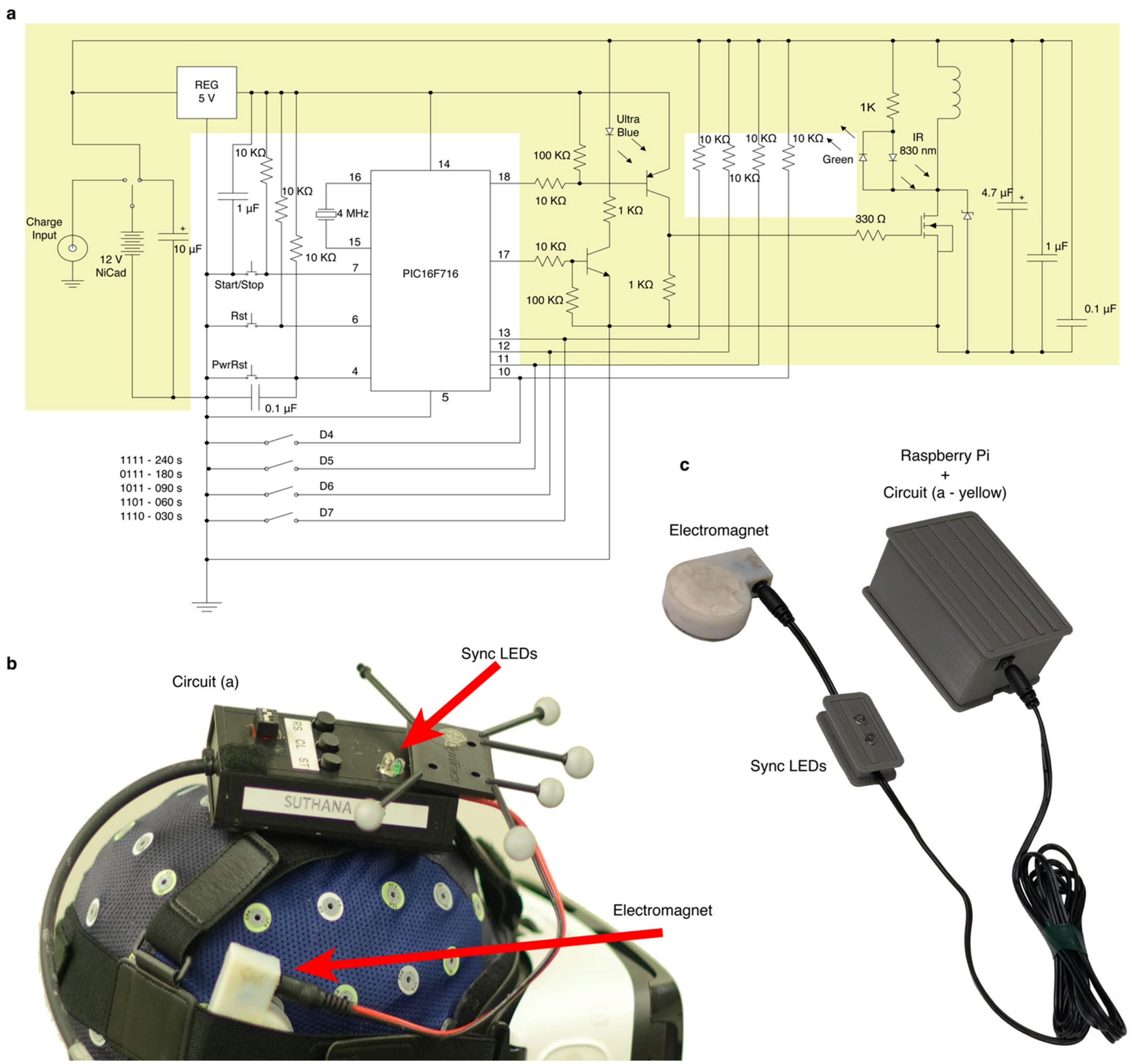


**Fig. S2 | Programmable electromagnet device for Mo-DBRS Lite. a**, Schematic of the electrical circuit necessary for driving an electromagnet, whose pulse is sent towards the underlying implanted RNS Neurostimulator. The system includes power delivery from a 12 V battery, various capacitors for voltage stabilization, current limiting resistors, and LEDs that emit light when the *Magnet* pulse is being sent that can be used for synchronization with external cameras. The electromagnet device can be manually controlled via button presses captured by a PIC controller that can deliver pulses every 30, 60, 90, 180, or 240 s (configurable by manually setting a 4-pin DIP switch). Such a device requires the complete circuitry shown in **a**, with the actual device worn by the participant shown in **b**. **c**, For wireless remote control of the electromagnet device, additional circuitry (marked in yellow **a**) can be added around a Raspberry Pi (RP) with pins 17 and 18 from a PIC controller being interfaced with GPIOs. The modified RP is shown in figure **c**, and can be remotely used to trigger the electromagnet device repeatedly with programmable times and durations from within the RP or the Experimental Task Program.

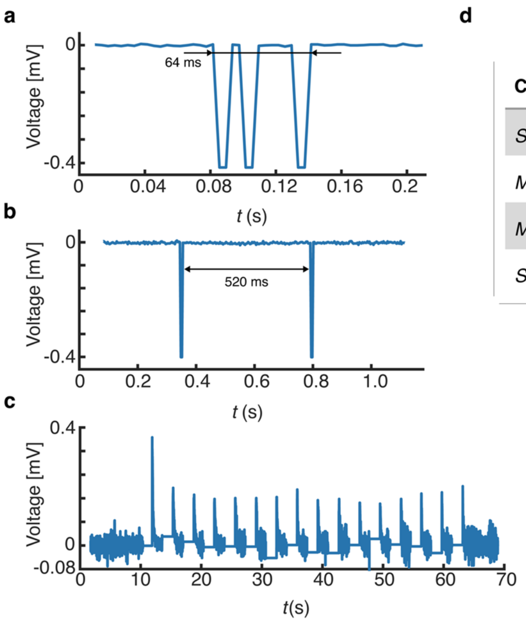


d

| Command | Tethered (Laptop) | Wireless (Laptop) | Wireless (Tablet) |
|---|---|---|---|
| *Store* | 3094.11 (45.69) | 3092.02 (46.13) | 1145.23 (35.87) |
| *Mark* | 1.51 (0.05) | 3.71 (0.15) | 2.65 (0.11) |
| *Magnet* | 13.54 (5.94) | 14.01 (7.13) | 13.23 (6.74) |
| *Stimulation* | 1415.61 (87.04) | 1419.78 (93.41) | 1355.51 (26.5) |

**All times in ms*

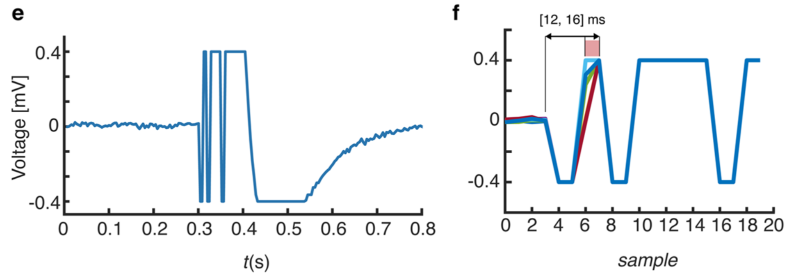


**Fig. S3 | Characterization of Mo-DBRS command latencies. a**, Example of a Real-Time iEEG recording during a remotely delivered *Mark* command tested in-vivo with an example participant. A single *Mark* signal has a distinctive pattern with 3 spikes, 16 sample points long (64 ms), and can be easily detected in post-processing procedures for synchronization purposes. If there is an insecure connection between the Wand and head that causes any displacement, then the complete *Mark* signal pattern is not captured, missing one or two spikes. Only the *Mark* signals that have at least 2 spikes can be used for accurate synchronization. **b**, Example of a remotely delivered *Magnet* signal recorded in Real-Time iEEG in-vivo. Shown is the full electromagnetic pulse with a 520 ms duration. **c**, In-vivo Real-Time iEEG activity showing 16 remotely delivered stimulation bursts (via the *Stim* command) with a 2.2 s inter-stimulation interval. **d**, Summarized average (error) latencies for each command (*Store*, *Mark*, *Magnet*, and *Stim*) from the Raspberry Pi (RP) during tethered and wireless setups. Note that the *Store* and *Stim* command latencies were measured with respect to the end of delivery while *Mark* and *Magnet* latencies were measured with respect to the first spike in the signal. **e**, Ex-vivo testing of the *Mark* command where shown is the Real-Time iEEG that captured a *Mark* signal along with a simultaneously sent RP test pulse that was sent to the same iEEG channel being recorded. **f**, Zoomed in Real-Time iEEG (from **e**), showing that the *Mark* signal arrives before the RP test pulse event. In 10 test trials shown here (different line colors), the offset was 12 to 16 ms between the *Mark* command and the RP test pulse. Given the sampling period of 4 ms, this thus reflects the synchronization latency that can be achieved using the *Mark* command.

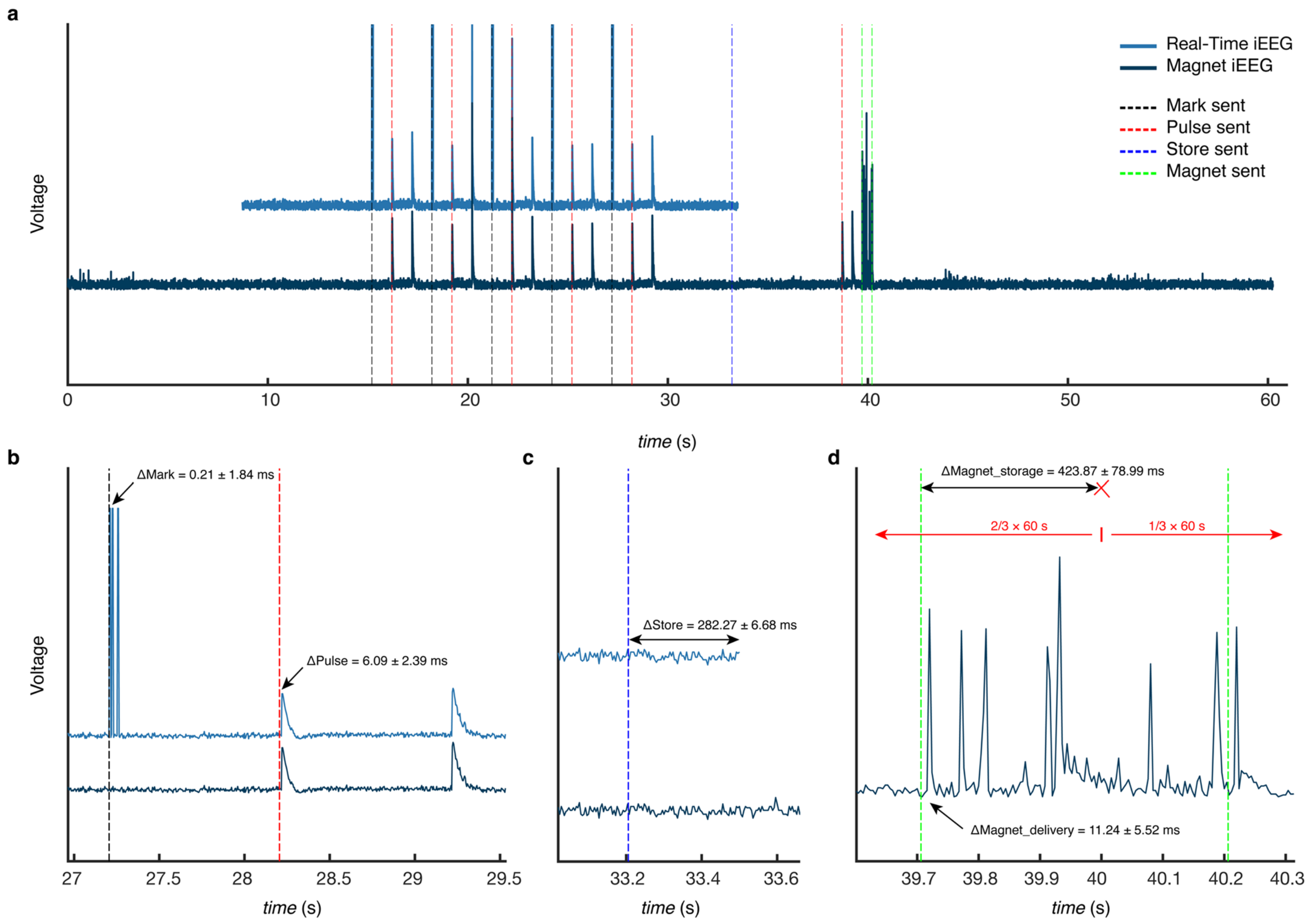


**Fig. S4 | Ex-vivo characterization of variability in Mo-DBRS Research Tool commands.** Variability in the *Mark*, *Store*, and *Magnet* commands was tested by sending a test signal (Pulse) from the Raspberry Pi (RP) to a test RNS System electrode contact. **a**, Shown is the simultaneously captured Real-Time iEEG and Magnet-triggered iEEG (absolute values) with a sequence of commands sent: 5 × (*Mark*, Pulse), *Store*, Pulse, *Magnet* (pre-configured in the Programmer to capture iEEG with a duration of 60 s). Dotted lines represent timestamps when the corresponding event was initiated on the RP. **b**, Zoomed in portion showing a *Mark* signal and Pulse signal where the uncertainty of delivery was determined to be 0.21 and 6.09 ms respectively. **c**, Captured relative offset between the *Store* command initiation and actual stored Real-Time iEEG where variability was measured to be 282.27 ms. **d**, Variability in the *Magnet* command delivery was determined to be 11.24 ms whereas storage variability of the Magnet-triggered iEEG data was determined to be 423 ms. Note that 40 s here (red cross mark) is the point where 2/3 (1/3) of the Magnet-triggered iEEG data that was stored occurred before (after).

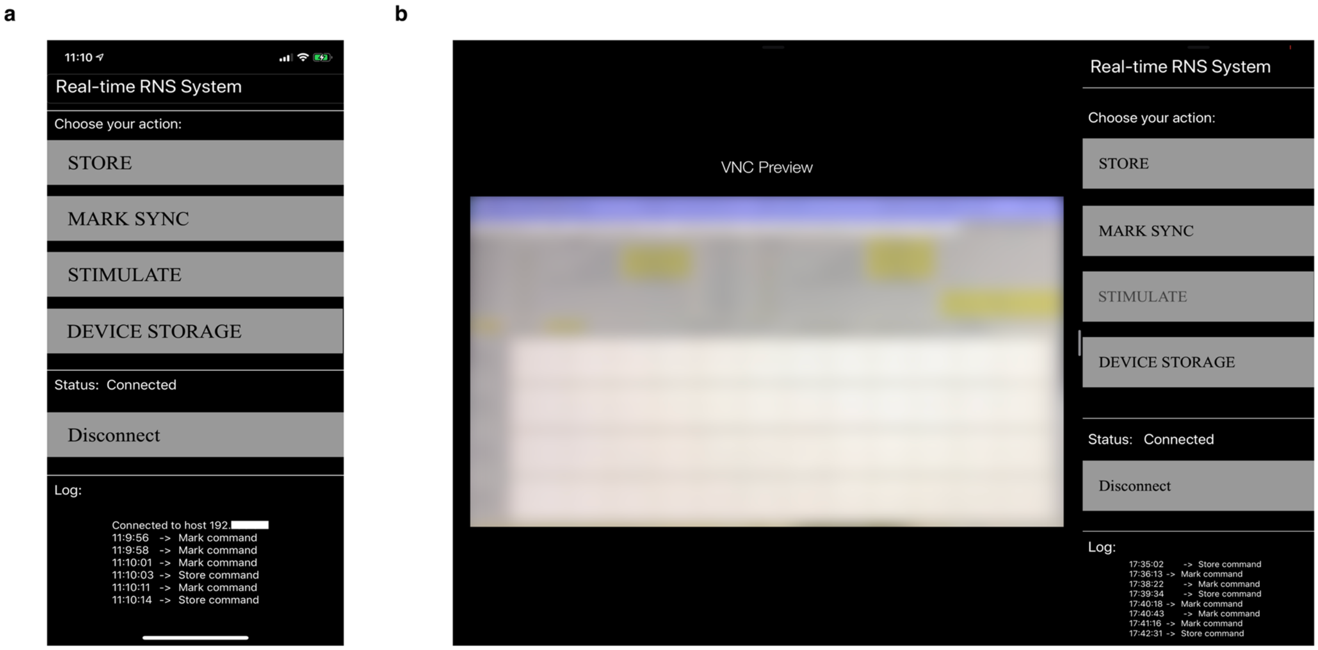


**Fig. S5 | Manual remote command interface developed for iOS devices. a,** iPhone app**. b,** iPad app provides more space for viewing the Programmer VNC connection. Apps were written in Swift 5 programming language using Xcode 10 Environment and are available on GitHub.

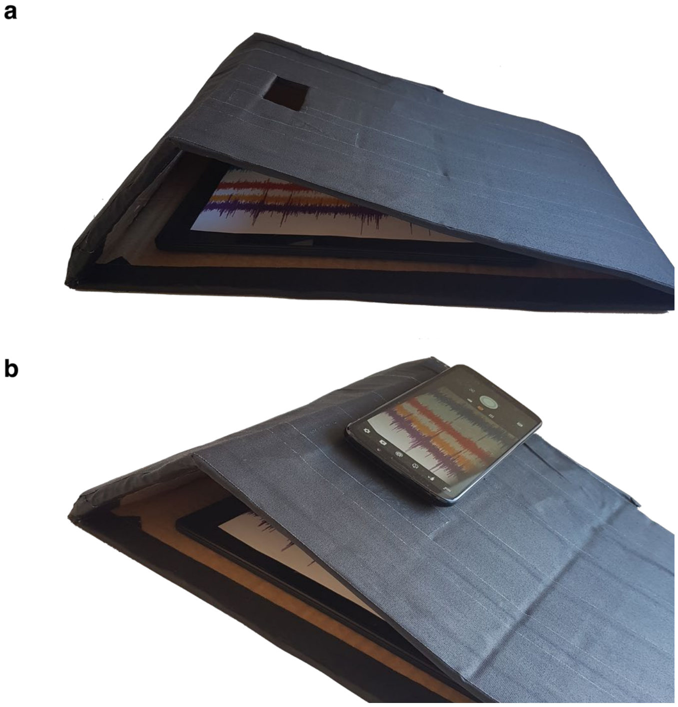


**Fig. S6 | Indirect real-time remote iEEG viewing for the Tablet Programmer.** The Laptop Programmer allows remote access/viewing, but the Tablet Programmer does not. **a**, To indirectly and remotely view Real-Time iEEG, the Tablet Programmer can be enclosed in a custom-shaped box and **b**, a phone camera can be used to record the Tablet's screen and stream the video to the Experimental Computer via TeamViewer.

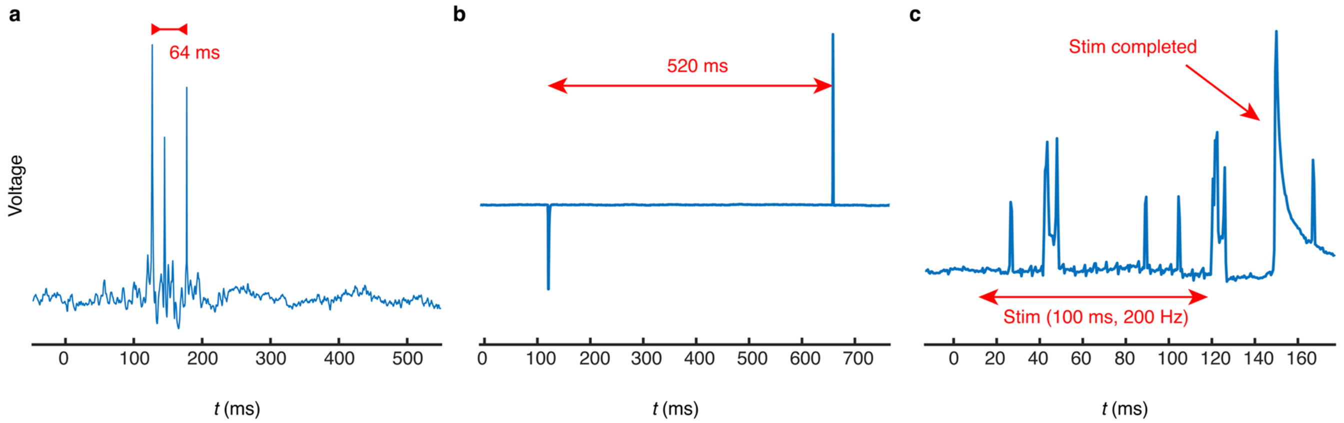


**Fig. S7 | *Mark*, *Magnet*, and *Stim* signals recorded in scalp EEG that can be used for synchronization with iEEG . a**, Example scalp EEG captured *Mark* signal that can be aligned with Real-Time iEEG. **b**, Example scalp EEG captured *Magnet* signal that can be aligned with Real-Time iEEG (*Magnet* artifact) and Magnet-triggered iEEG (logged timestamp). **c**, Example scalp EEG captured *Stim* signal that can be aligned with Real-Time iEEG and Magnet-triggered iEEG.

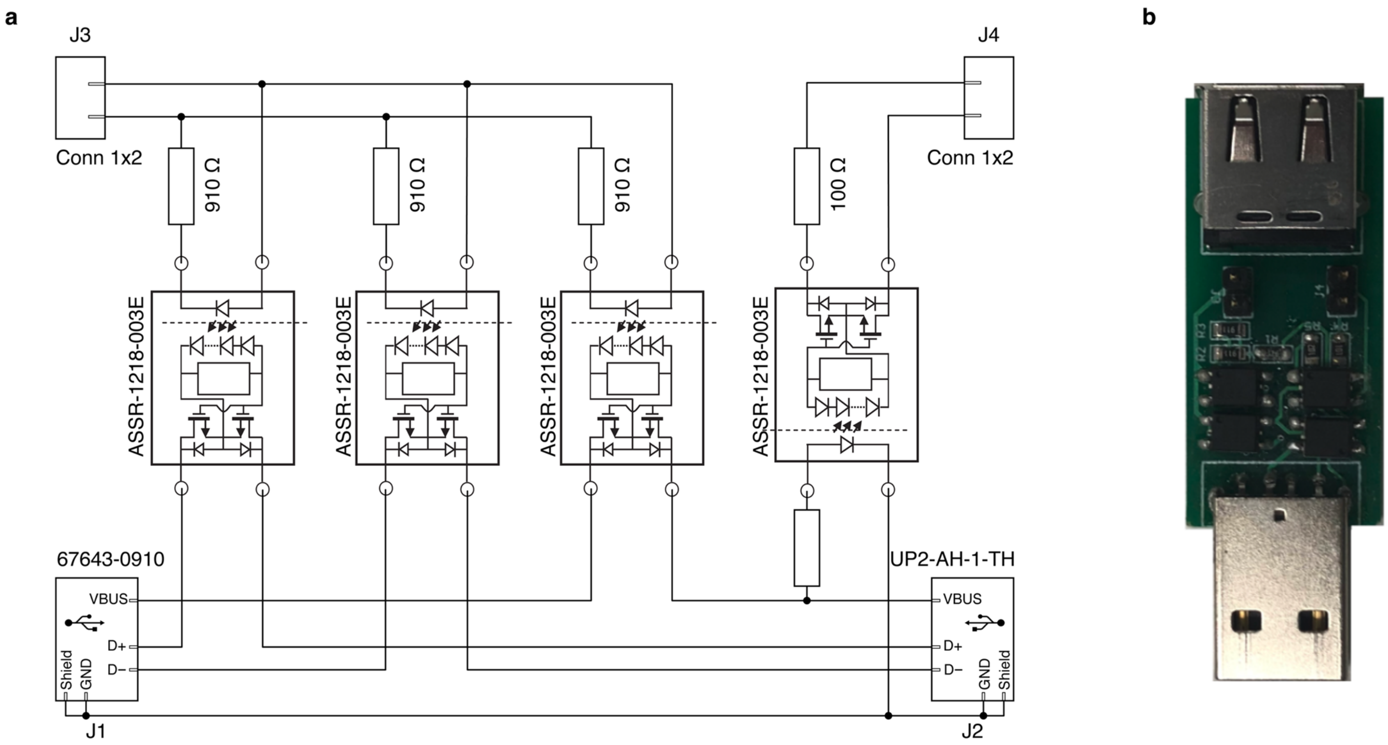


**Fig. S8 | Telemetry Switch. a**, Schematic of the USB switch circuit used for turning on/off telemetry. In the center, there are four optical switches attached to the USB power and data lines. An optical switch on the far right is used to read the current status of the USB power lines. A J3 connector is connected to the Raspberry Pi GPIO output and serves as a digital control of the switch state. The J4 connector is connected to the Raspberry Pi GPIO input and provides the status of the communication. J1/J2 are standard USB female/male connectors **b**, The actual USB switch device that was manufactured.

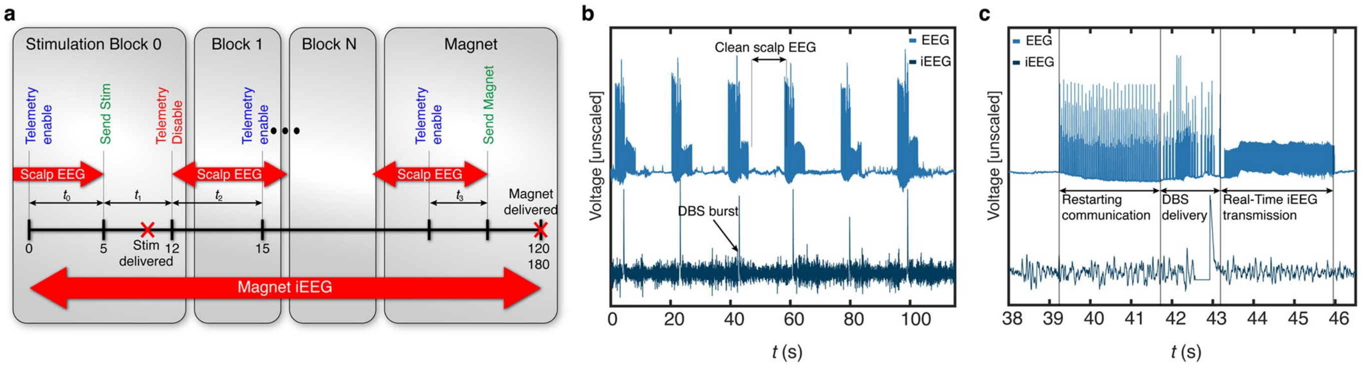


**Fig. S9 | Procedure for using the Telemetry Switch solution for simultaneous iEEG + scalp EEG. a**, Proposed procedure for delivering a series of DBS bursts with minimally contaminated scalp EEG activity. Timings $t_0$, $t_1$, $t_2$, $t_3$ denote the telemetry enable time, command delivery time, clean scalp EEG recording time, storing of Magnet-triggered iEEG time after enabling communication, respectively. Note that scalp EEG recording is clean during $t_0$ until communication is established again. **b**, Magnet-triggered iEEG and scalp EEG recordings were acquired following the **a** procedure involving 6 DBS bursts of 100 ms duration. Cycles of clean scalp EEG and recordings containing telemetry artifacts are visible. iEEG/EEG recordings were synchronized by aligning DBS artifacts in both recordings. **c**, One cycle of synchronized parallel recording of Magnet-triggered iEEG and scalp EEG from **b**. Three different periods of telemetry artifacts (Restarting of communication, DBS, and Real-Time iEEG) are denoted.

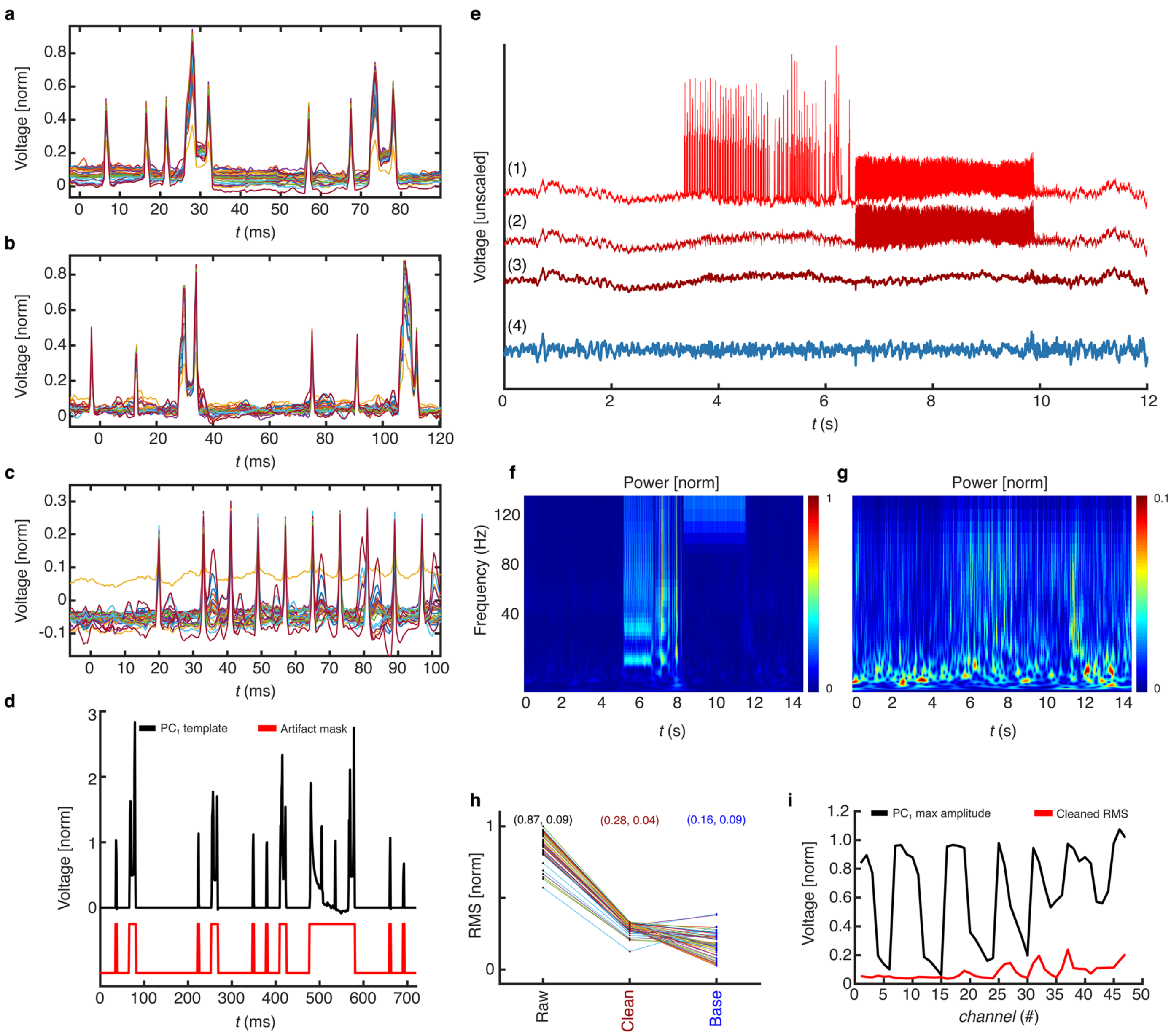


**Fig. S10 | Offline artifact rejection solution for simultaneous iEEG and scalp EEG. a**, Close look at each of the telemetry noise patterns recorded from scalp EEG channels is plotted on **a** – **c**. **a**, Zoomed-in pattern of the scalp EEG artifacts related to the restarting of telemetry is represented as a repetitive sequence of the shown waveform. **b**, Zoomed-in pattern of scalp EEG artifacts characteristic of DBS delivery is represented as a repetitive sequence of the shown waveform. **c**, Zoomed-in pattern of scalp EEG artifacts characteristic for Real-Time iEEG transmission is represented as a 125 Hz sequence of spikes. **d**, PCA analysis of the scalp EEG data across all channels captures most of the shared artifacts in the first PC1 component (solid black line). Information on timings and shapes of each part of the artifact is used to extract the exact position of segment affected by telemetry. An artifact mask (solid red line) is used to separate artifact spikes from PCA components and discard the rest of the picked-up correlation between scalp EEG channels. **e**, The three steps used for artifact reduction shown in one example scalp EEG channel: 1. Subtraction of the first 3 PC components from the scalp EEG; 2. Downsampling to 250 Hz; 3. Reducing outstanding (as per base scalp EEG statistics) spikes in the Wavelet domain and applying a 2 Hz high pass filter. 4. Resulting cleaned scalp EEG channel after artifact reduction steps 1-3. **f**, Time-frequency power scaleogram of raw scalp EEG. **g**, Time-frequency power scaleogram of cleaned scalp EEG. **h**, Normalized RMS value of the signal, with mean and standard deviation values, for all channels before and after cleaning, as well as comparison with RMS of the corresponding base scalp EEG, unaffected by the artifacts. **i**, Normalized amplitudes of extracted PC1 component across channels describes the distance between the Wand and scalp EEG channels (solid black line). The RMS value of cleaned scalp EEG across channels (solid red line) shows that the results of the algorithm slightly deteriorate in the immediate vicinity to the Wand, where telemetry is best picked up by those channels (e.g., 32 – 47).

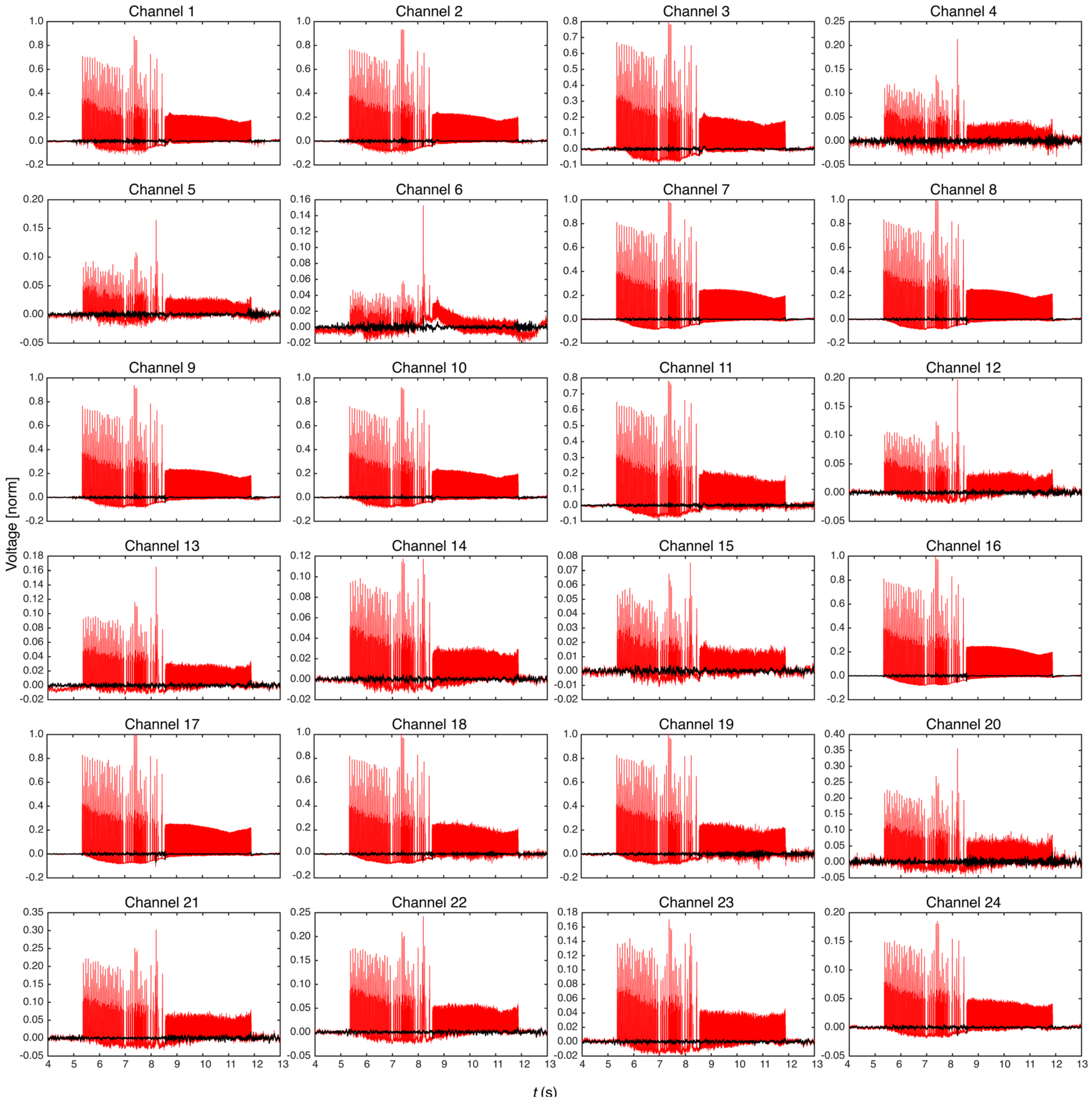


**Fig. S11 | Additional scalp EEG channels (1 – 24) before (red) and after (black) cleaning.**

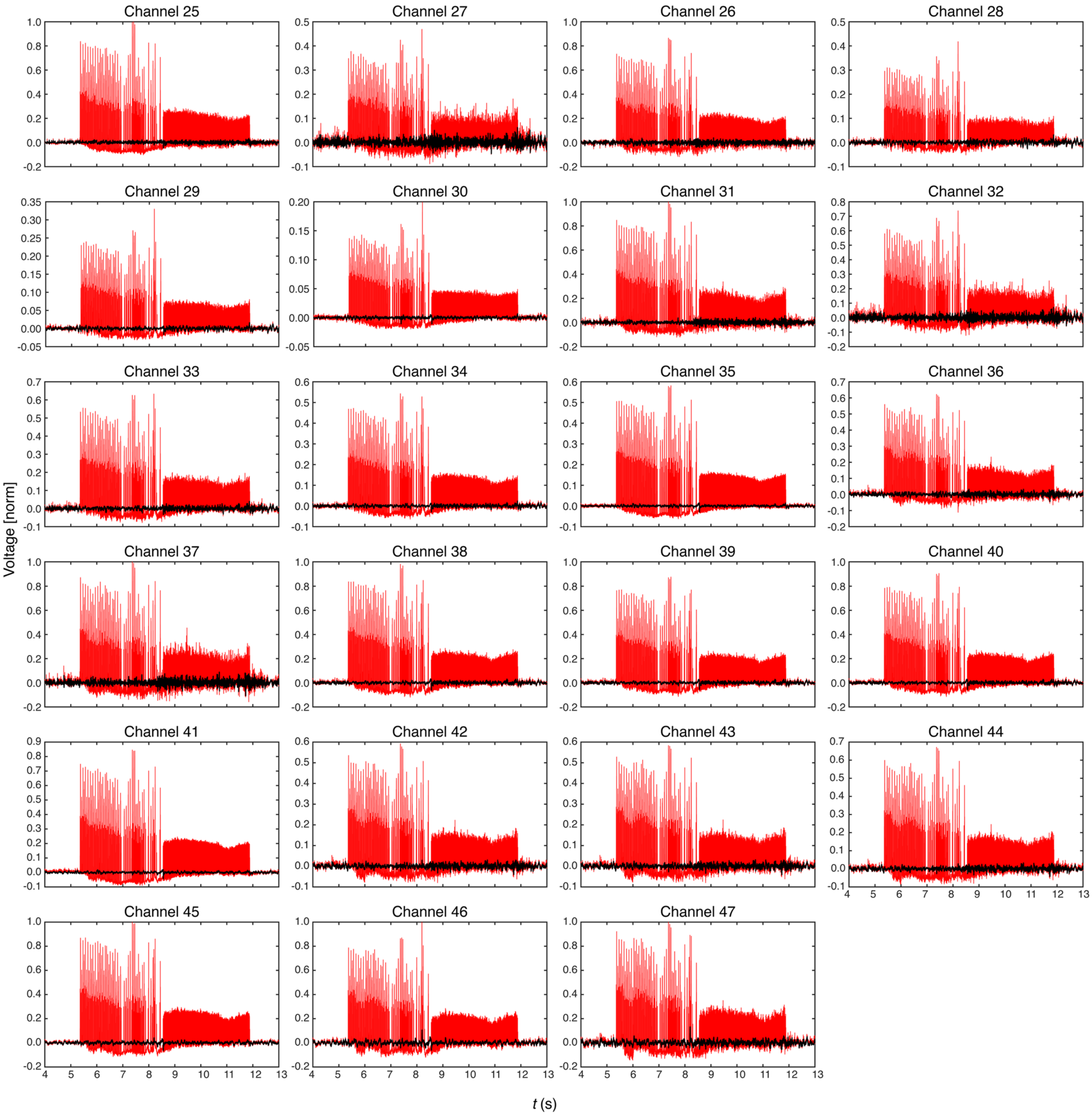


**Fig. S12 | Additional scalp EEG channels (25 – 47) before (red) and after (black) cleaning.**

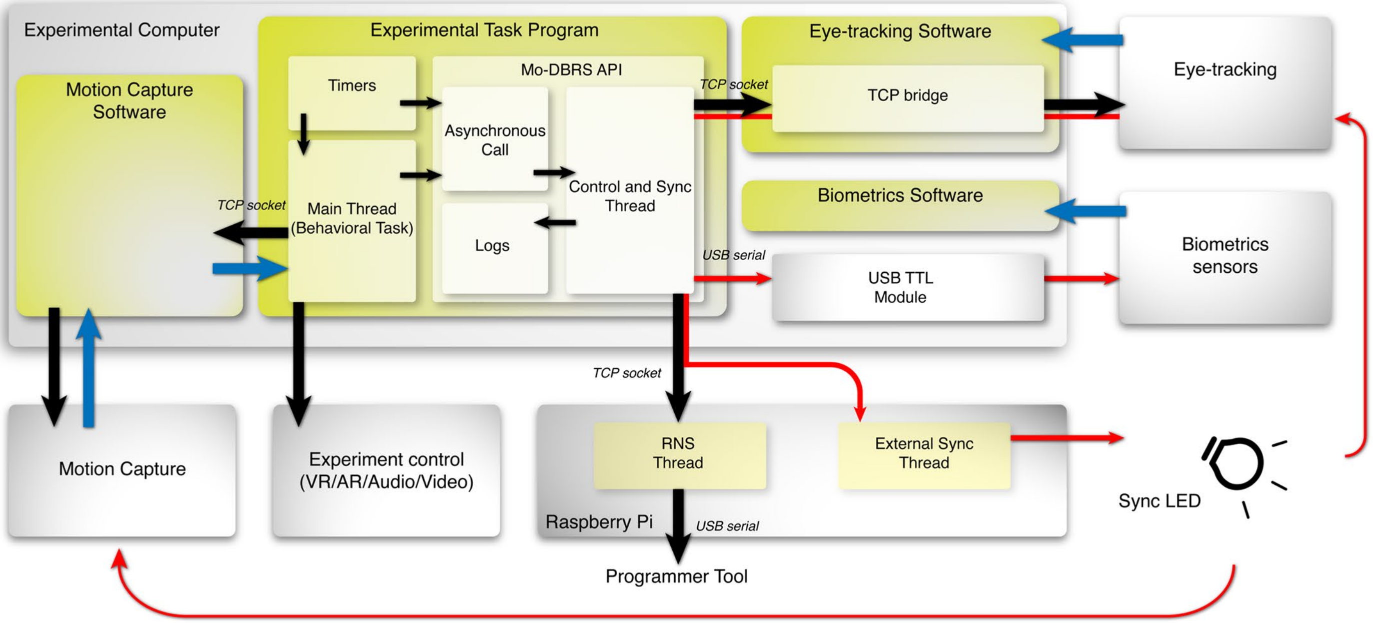


**Fig. S13 | Block diagram of an example Experimental Task Program.** Here we present the general block diagram for a suggested Experimental Task Program that can be adapted for any of the commonly used software environments (e.g., Unity, Python, Matlab). Black arrows show control/configuration flow, blue arrows show data flow, and red arrows point to synchronization between different components. **Main Thread** contains the software running the behavioral task, any online processing of behavior that may be used to update the VR/AR/Audio/Video stimuli (Experimental control). **Timers** keep track of the task and provide necessary time information to the Main Thread and Sync Thread. Although usually part of the Main Thread, timers are separated here in case RNS commands need to be sent independently of the Main Thread (Synchronous). Synchronous events are triggered periodically from the timer block, while asynchronous events can be called directly from the Main Thread. **Asynchronous Call** block contains the available functions that are sent to target devices (e.g., RNS and other wearable sensors) which are part of the proposed **Mo-DBRS API**. **Control and Sync Thread** synchronously (periodically) or asynchronously (on-demand) sends the commands to the targeted device, as well as broadcasts simultaneous synchronization signals to all connected wearable sensors. All traffic timestamps are being logged into the **Logs** local storage. As mentioned previously, the Experimental Task Program, motion capture system (e.g., Optitrack), implanted RNS System, and eye-tracking (e.g., Pupil Labs) are communicating through the local network, while biometric measurements (e.g., Biopac) are using a USB connection and external USB Module for synchronization. The Raspberry Pi contains a **RNS Thread** running the server, which accepts incoming commands from the Experimental Task Program. To bypass pipeline delays, an **External Sync Thread** turns on an external LED for 50 ms through GPIO whenever the *Mark* (*Magnet*) command is received on the Raspberry Pi. An emitted LED is captured by wall-mounted (e.g., OptiTrack) or wearable cameras, and/or eye-tracking (e.g., Pupil Lab) cameras, thus providing additional synchronization.

| Participant | Age | Gender | Handedness |
|---|---|---|---|
| $P_1$ | 21 | M | R |
| $P_2$ | 39 | F | L |
| $P_3$ | 40 | M | R |
| $P_4$ | 48 | F | R |
| $P_5$* | 65 | M | R |

**Table S1 | Participant Demographics.** Demographics of the study participants including age, gender and handedness.
* Participant P5 is congenitally blind.

| Participant | Contact Labels | Electrode Spacing | Contact 1 | Contact 2 |
|---|---|---|---|---|
| $P_1$ | RHIP1 – RHIP2 | 3.5 mm | -- | Subiculum |
| | RHIP3 – RHIP4 | | -- | CA1 |
| | ROFC1 – ROFC2 | | Orbitofrontal Cortex | Orbitofrontal Cortex |
| | ROFC3 – ROFC4 | 10 mm | Orbitofrontal Cortex | Orbitofrontal Cortex |
| $P_2$ | LHIP1 – LHIP2 | 10 mm | CA1 | CA1 |
| | LHIP3 – LHIP4 | | CA1 | CA1 |
| | REC1 – REC2 | | Entorhinal Cortex | Perirhinal Cortex |
| | REC3 – REC4 | 3.5 mm | Perirhinal Cortex | Fusiform Gyrus |
| $P_3$ | LAH1 – LAH2 | 3.5 mm | CA23DG | CA23DG |
| | LAH3 – LAH4 | | CA23DG | CA23DG |
| | RAH1 – RAH2 | | Subiculum | Subiculum |
| | RAH3 – RAH4 | 3.5 mm | CA23DG | CA1 |
| $P_4$ | LEC1 – LEC2 | 10 mm | Hippocampus | Perirhinal Cortex |
| | LEC3 – LEC4 | | Inferior Temporal | Inferior Temporal |
| | RAH1 – RAH2 | | Subiculum | Subiculum/CA1 |
| | RAH3 – RAH4 | 3.5 mm | CA1 | Perirhinal Cortex |
| $P_5$ | LEC1 – LEC2 | 10 mm | Entorhinal Cortex | Perirhinal Cortex |
| | LEC3 – LEC4 | | Inferior Temporal | Inferior Temporal |
| | LHIP1 – LHIP2 | | CA1 | CA1 |
| | LHIP3 – LHIP4 | 3.5 mm | CA1 | CA1 |

**Table S2 | Electrode Localization.** Electrode contact locations from the participants in the study. Bipolar recordings were performed between Contact 1 and 2 (3 and 4). Locations were targeted for electrode placement based on clinical criteria and include R(L)EC, corresponding to the right (left) entorhinal cortex, R(L)HIP corresponding to the right (left) hippocampus, ROFC corresponding to the right orbitofrontal cortex, R(L)AH corresponding to the right (left) anterior hippocampus.

| Setup Options | Real-Time View | Programmable DBS | Task - iEEG Synchronization | Offset (ms) | Scalp EEG - iEEG Synchronization | Offset (ms) | Telemetry artifact solution for scalp EEG | Wearability |
|---|---|---|---|---|---|---|---|---|
| Mo-DBRS<br>Real-Time iEEG | ✓ | ✓ | *Mark* | 14 ± 2 | *Mark* | 16 ± 4 | Artifact rejection | Full wearable platform |
| Mo-DBRS Lite<br>Magnet iEEG | ✗ | ✗ | *LED* | 432 ± 83 | *Magnet* | 424 ± 79 | None | Lightweight wearable platform |
| Mo-DBRS<br>Real-Time +<br>Magnet iEEG | ✓ | ✓ | *Mark/ Magnet* | 14 ± 2 /<br>424 ± 79 | *Mark / Magnet / DBS* | 16 ± 4 /<br>426 ± 81 /<br>2 ± 2 | Telemetry switch<br>+<br>Artifact rejection | Full wearable platform |

**Table S3 | Considerations that can guide selection of platform version (Mo-DBRS or Mo-DBRS Lite).** Shown are the setup options and whether they allow for Real-Time iEEG, Magnet-triggered iEEG (Magnet iEEG) or deep brain stimulation (DBS). Also shown are the methods available for synchronization of iEEG with the Experimental Task Program (task-iEEG) and scalp EEG (Scalp-iEEG synchronization), the associated latency offsets and solutions for minimization of telemetry related artifacts. Presented offsets are rounded and in the format of mean ± std.
